# Transcriptional and spatial profiling of fibroblasts from human lungs highlights CTHRC1+ cells as fibrogenic signaling hubs in fibrosis

**DOI:** 10.64898/2026.04.08.717092

**Authors:** N.D. Vanegas-Avendano, H. Chen, J. Wellmerling, J. Rodriguez-Lopez, A. Ghobashi, V. Peters, C. Sen, B. F. Reader, K. Shilo, B. Gomperts, Q. Ma, A.L. Mora, D.J. Tschumperlin, M. Rojas

**Affiliations:** Division of Pulmonary, Critical Care, & Sleep Medicine, Department of Internal Medicine, The Ohio State University, Wexner Medical Center, Columbus, OH, USA; Department of Biomedical Informatics, College of Medicine, The Ohio State University, Columbus, OH, USA; Pelotonia Institute for Immuno-Oncology, The James Comprehensive Cancer Center, The Ohio State University, Columbus, OH, USA; Department of Pathology, The Ohio State University Wexner Medical Center, Columbus, OH, USA; Human Tissue Biorepository. OSU Wexner Medical Center Comprehensive Transplant Center, The Ohio State University, Columbus, OH, USA; Department of Physiology & Biomedical Engineering. Mayo Clinic College of Medicine and Science, Rochester, Minnesota, USA; Children’s Discovery and Innovation Institute, Broad StemCell Research Center, Jonsson Comprehensive Cancer Center, Departments of Pediatrics and Pulmonary Medicine, University of California, Los Angeles, California, USA

## Abstract

Lung fibroblasts are key regulators of tissue homeostasis and extracellular matrix (ECM) remodeling, and their aberrant activation drives the progressive parenchymal scarring characteristic of idiopathic pulmonary fibrosis (IPF), a fatal disease with limited therapeutic options. Despite their central pathogenic role, lung fibroblasts are difficult to isolate due to their embedded position within the ECM, and standard in vitro culture conditions may lead to the loss of their native functional and transcriptional characteristics, hampering the study of fibroblast behavior in disease. The transcriptional heterogeneity of lung fibroblast subtypes and the extent to which culture-induced alterations diverge from native tissue signatures remain poorly understood. Here, we integrated single-cell RNA sequencing (scRNA-seq) and spatial transcriptomics of lung tissue from IPF patients and age-matched healthy donors with transcriptomic profiling of cultured fibroblasts collected at passages 1 and 6 after isolation using three optimized protocols: whole lung cell suspension (WLCS), negative fraction enrichment, and outgrowth. Tissue-based analysis identified six transcriptionally distinct mesenchymal subtypes: alveolar, adventitial, inflammatory, peribronchial, CTHRC1+ and smooth muscle cell (SMC). The fibroblast subtype CTHRC1+ represented the most transcriptionally activated pro-fibrotic subtype, showing the greatest upregulation of ECM biosynthesis genes, a prominent role in intercellular communication, and preferential enrichment within fibroblastic foci in IPF lung tissue. Pseudotime trajectory analysis supported a directional transcriptional continuum from alveolar and inflammatory fibroblasts toward the CTHRC1+ state, driven by coordinated activation of pro-fibrotic transcription factors, including RUNX2, CREB3L1, and SCX. In vitro culture progressively reshaped fibroblast transcriptional identity relative to native tissue, with increased collagen and matrix metalloproteinase (MMP) expression during passaging, loss of distinct CTHRC1+ fibroblasts, and gain of alveolar fibroblasts displaying pro-fibrotic activation across all isolation protocols. These findings provide a high-resolution transcriptional map of lung fibroblast heterogeneity in IPF and highlight critical limitations of standard in vitro culture systems for recapitulating native fibroblast diversity, with important implications for the development and evaluation of fibroblast-targeted therapeutic strategies in IPF.

## Introduction

Pulmonary fibrosis is a devastating lung disease characterized by excessive collagen deposition and aberrant extracellular matrix (ECM) remodeling, leading to progressive loss of lung function and, ultimately, respiratory failure. Idiopathic pulmonary fibrosis (IPF), the most common and severe form of pulmonary fibrosis, is associated with poor prognosis and limited therapeutic options. Fibroblasts are key cellular mediators of ECM synthesis, remodeling, and maintenance, and play a central role in both physiological lung homeostasis and pathological fibrotic processes.^1^ Consequently, understanding the mechanisms regulating fibroblast activation and collagen production is critical for the development of effective antifibrotic therapies for IPF.

In vitro culture systems have emerged as powerful tools for investigating lung fibroblast biology. These systems allow researchers to isolate and manipulate fibroblasts under controlled conditions, enabling detailed analyses of gene expression, protein production, and responses to experimental stimuli. However, a significant challenge associated with in vitro studies is the potential for culture-induced alterations in fibroblast phenotype.

In their native lung environment, fibroblasts reside within a complex three-dimensional (3D) architecture and interact with a diverse array of neighboring cell types, including immune, endothelial, and epithelial cells. This intricate cellular crosstalk, together with the biochemical and structural properties of the surrounding ECM, significantly influences fibroblast behavior.^2^

Isolation and subsequent in vitro culture on rigid plastic surfaces in a two-dimensional (2D) environment can disrupt these crucial interactions, leading to phenotypic changes that may alter fibroblast transcriptional and functional states. For instance, studies have shown that in vitro culture can result in the loss of differentiated fibroblast markers and activation toward a contractile myofibroblast phenotype.^3–5^ These observations underscore the need for optimized isolation and culture approaches that minimize phenotypic drift and better preserve the in vivo characteristics of lung fibroblasts. While in vitro 2D culture provides a controlled environment for genetic manipulation and remains a widely used and accessible experimental platform, it is important to recognize its limitations in fully recapitulating the in vivo microenvironment.

Integrating improved isolation and culture approaches with single-cell RNA-sequencing (scRNA-seq) provides an opportunity to more comprehensively characterize fibroblast heterogeneity and the transcriptional programs driving pulmonary fibrosis. To address these challenges, we investigated the transcriptional heterogeneity of human lung fibroblasts in both native lung tissue and in vitro culture using an integrative single-cell sequencing and spatial transcriptomic approach. By comparing fibroblast-enriched populations isolated from IPF patients and age-matched healthy donors using three optimized isolation protocols, whole lung cell suspension (WLCS), negative fraction enrichment, and outgrowth, we systematically evaluated how isolation methods and serial passaging influence fibroblast subtype composition and gene expression relative to native tissue.

Our study reveals how in vitro culture systematically reshapes ECM-related transcriptional programs, particularly collagen and matrix metalloproteinase (MMP) expression, and drives the selective expansion or loss of specific fibroblast subpopulations. These changes include alveolar fibroblast populations implicated in lung fibrosis^6–8^, as well as CTHRC1+ fibroblasts, which we identify as the most transcriptionally activated pro-fibrotic subtype in IPF tissue.

By bridging the gap between in vivo fibroblast heterogeneity and culture-induced transcriptional dynamics, this study advances our understanding of how in vitro conditions reshape fibroblast identity and function, and how these changes may influence the interpretation of in vitro experimental results. Together, these findings provide a framework for developing more representative in vitro models and for designing therapeutic strategies that target specific fibroblast subtypes and their distinct contributions to IPF pathophysiology.

## Results

### Fibroblast heterogeneity and spatial distribution across IPF and aged healthy lungs

To characterize the cellular landscape of fibroblast heterogeneity in IPF, we performed a multi-capture scRNA-seq study using lung tissue collected from age-matched healthy donors (>55 years; n = 10) and IPF explants sampled from both upper and lower lobes (n = 7 lungs/group; 14 IPF lobes). Tissues were processed into formalin-fixed paraffin-embedded (FFPE) blocks and sectioned for spatial transcriptomic analyses (**Fig. 1a**). After quality filtering and doublet removal, a total of 24 samples were retained for downstream analysis. Unsupervised clustering and uniform manifold approximation and projection (UMAP) visualization of all recovered cells identified major lung cell lineages, including epithelial, endothelial, immune, and mesenchymal compartments, with clear separation of clusters by lineage (**Fig. 1b**). Lineage identity was confirmed by expression of canonical marker genes, with mesenchymal and fibroblast cell populations defined by markers including *COL1A1* and *PDGFRA* (**Fig. 1c**). Heatmap analysis further confirmed robust and specific enrichment of lineage markers across major cell populations (**Fig. 1c**). Cell composition analysis across sample groups revealed an increased proportion of mesenchymal cells in IPF samples compared with age-matched control samples, consistent with the fibroproliferative pathology characteristic of this disease (**Fig. 1d**).

**Figure 1.**
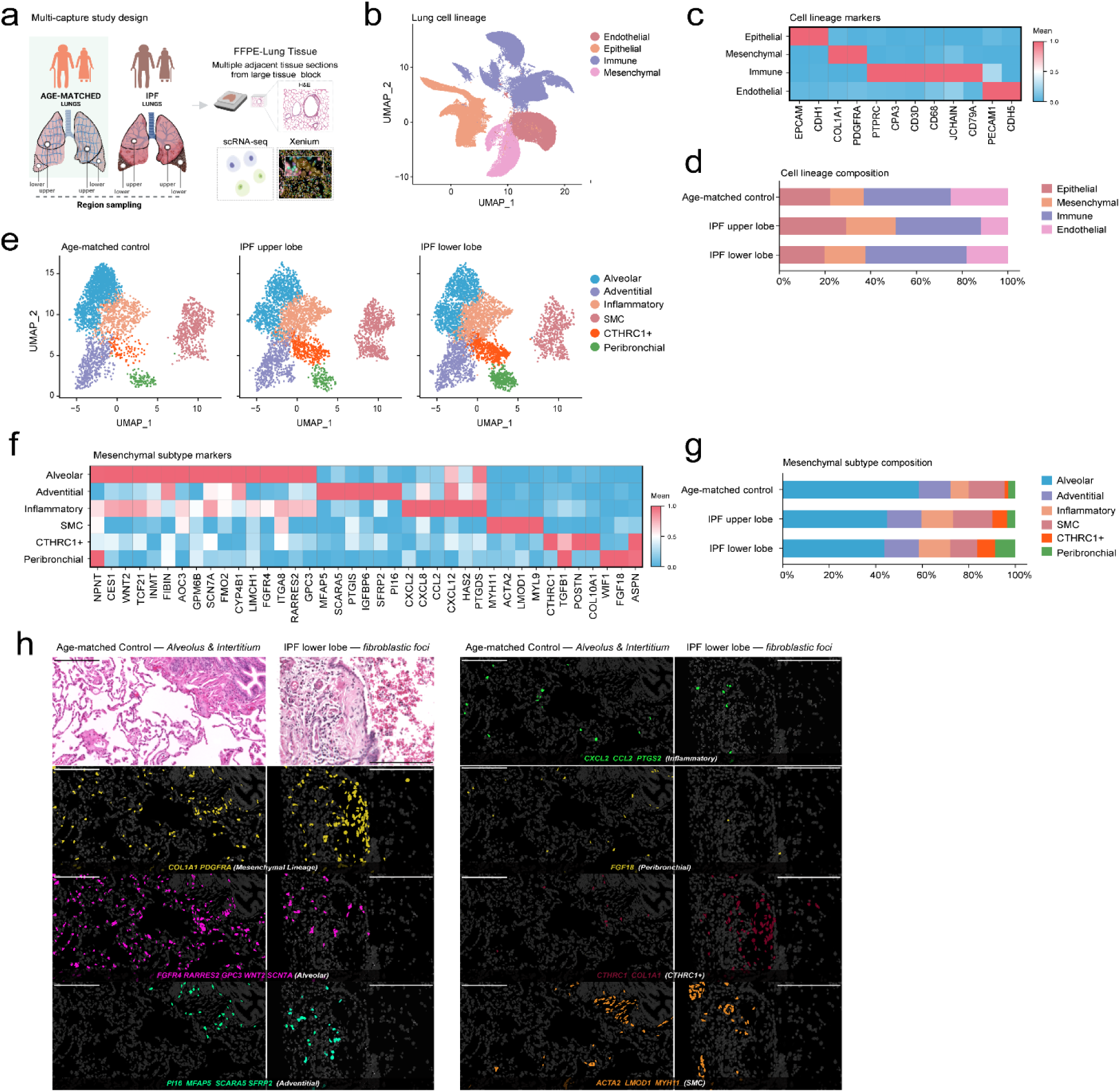
Single-cell analysis of mesenchymal heterogeneity and spatial distribution in IPF and aged healthy lungs. **a.** Schematic of the multi-capture study design, including lung tissue collection from age-matched healthy donors and IPF explants, preparation of FFPE sections, and downstream scRNA-seq and Xenium spatial transcriptomic analyses. **b.** UMAP visualization of major lung cell lineages across 24 samples, including age-matched controls and IPF upper and lower lobes. **c.** Heatmap of lineage-specific marker gene expression. **d.** Cell composition analysis showing an increased mesenchymal cell proportion in IPF samples. **e.** Secondary clustering of the mesenchymal compartment across age-matched control samples and IPF upper and lower lobe samples; UMAP representation of the six mesenchymal subtypes. **f.** Heatmap of marker gene expression for the six mesenchymal subtypes after sample integration. **g.** Distribution of mesenchymal subtypes by sample group (age-matched control, IPF upper lobe, and IPF lower lobe). **h.** Spatial distribution of mesenchymal subtypes in lung samples. Representative H&E micrographs and Xenium maps of selected mesenchymal subtype markers in age-matched control alveolar/interstitial regions and IPF lower lobe fibroblastic foci. H&E micrographs illustrating tissue architecture are shown at the top left, and gene markers used to identify each fibroblast subtype are indicated on the individual panels. Scale bar, 200 μm.

To resolve fibroblast subpopulation heterogeneity, we performed secondary clustering restricted to the mesenchymal compartment, which resolved six transcriptionally distinct fibroblast subtypes across age-matched control, IPF upper lobe, and IPF lower lobe samples (**Fig. 1e, Supplementary Fig. 1a**). These fibroblast subpopulations were designated adventitial, alveolar, inflammatory, CTHRC1^+^, peribronchial, and SMC based on their top differentially expressed marker genes. Heatmap analysis of subtype-specific marker gene expression confirmed transcriptional distinctiveness across all six populations, with alveolar marked by *NPNT*, *CES1*, *TCF21, INMT*, *FIBIN*, and *AOC3*; adventitial by *MFAP5, SCARA5, SFRP2* and *PI16*; and peribronchial by *WIF1*, *FGF18*, and *ASPN*. The inflammatory subtype was distinguished by a chemokine-rich signature (*CXCL2*, *CXCL8*, *CCL2*) and *HAS2*; CTHRC1+ fibroblasts by co-expression of *CTHRC1*, *TGFB1*, and *POSTN and* SMC by *MYH11*, *ACTA2*, *LMOD1*, and *MYL9* (**Fig. 1f**). Integrated fibroblast UMAP visualization colored by sample of origin showed broad intermixing of cells from age-matched control, IPF upper lobe, and IPF lower lobe samples across the major fibroblast clusters, supporting shared subtype structure across sample groups (**Supplementary Fig. 1a**). Pericytes were not included in the present fibroblast subtype analysis, as they represent a functionally and anatomically distinct mesenchymal population.^9,10^ Their perivascular confinement and vascular-supportive functions of pericytes place them outside the scope of the interstitial fibroblast biology addressed in this study.

Having established the transcriptional identity of each fibroblast subtype using canonical marker genes, we next examined how the expression of subtype-specific markers of all six fibroblast subtypes varies across age-matched control, IPF upper lobe, and IPF lower lobe samples. Psuedobulk analysis revealed distinct transcriptional profiles across fibroblast subtypes in IPF upper and lower lobes relative to age-matched healthy donors, with CTHRC1+ fibroblasts characterized by elevated expression of ECM-related genes, including *CTHRC1*, *COL10A1*, and *POSTN* (**Supplementary Fig. 1b**). This signature was most prominent in IPF samples, particularly IPF lower lobe samples, consistent with the increased disease severity typically observed in lower lung zones. In contrast, alveolar fibroblasts were defined by high expression of *INMT*, *FIBIN, AOC3, GPM6B, FGFR4,* and *RARRES2*; adventitial fibroblasts by *MFAP5, SCARA5, PI16*, and *SFRP2*; inflammatory fibroblasts by *CCL2*, *HAS2*, and *PTGDS*; peribronchial fibroblasts by *WIF1, FGF18,* and *ASPN*; SMCs by contractile markers such as *ACTA2*, *MYH11*, *LMOD1*, and *MYL9* (**Supplementary Fig. 1b**).

To validate the selective enrichment of *CTHRC1* and *COL1A1* in the CTHRC1+ subpopulation, we examined single-cell expression distributions across all fibroblast subtypes and groups (**Supplementary Fig. 1c, left**). Violin plots confirmed that *CTHRC1* expression was highly specific to the CTHRC1+ subpopulation, with minimal expression detected in adventitial, alveolar, inflammatory and peribronchial fibroblasts, or in SMCs across all groups (Supplementary Fig. 1c, left). *CTHRC1* expression was elevated in CTHRC1+ fibroblasts from IPF upper and IPF lower lobe samples relative to age-matched controls, with the highest levels observed in IPF lower lobe samples. Similarly, *COL1A1* exhibited the highest expression in CTHRC1+ fibroblasts, particularly in IPF samples, although lower-level expression was also detected in adventitial and peribronchial subtypes in the disease context (**Supplementary Fig. 1c, right**). The breadth and intensity of *COL1A1* expression in CTHRC1+ fibroblasts far exceeded that of other subtypes, highlighting their likely role as the primary collagen-producing cells in IPF lung tissue.

To directly compare *COL1A1* expression across fibroblast subtypes within each group, we stratified single-cell expression distributions by group (**Supplementary Fig. 1d**). In age-matched control samples, *COL1A1* expression was distributed at low levels across adventitial, alveolar, and peribronchial fibroblasts, with detectable expression also present in CTHRC1+ fibroblasts, although this population remained proportionally small in age-matched control samples (**Supplementary Fig. 1d, left; Fig. 1g**). In IPF upper lobe samples, CTHRC1+ fibroblasts emerged as the dominant *COL1A1*-expressing population, with significantly elevated expression relative to all other subtypes (**Supplementary Fig. 1d, middle**).

This pattern was further amplified in IPF lower lobe samples, where CTHRC1+ fibroblasts exhibited the highest *COL1A1* expression intensity and the broadest distribution, consistent with their expansion and activation in regions of advanced fibrosis (**Supplementary Fig. 1d, right**). Notably, inflammatory fibroblasts and SMCs maintained consistently low *COL1A1* expression across all groups, indicating that these subtypes likely do not substantially contribute to pathological collagen deposition in IPF lungs. Collectively, these findings establish that CTHRC1+ fibroblasts represent a transcriptionally distinct, disease-enriched subpopulation defined by coordinated upregulation of pro-fibrotic ECM genes, with *COL1A1* serving as a hallmark effector molecule driving fibrotic remodeling in IPF lung tissue. Quantification of fibroblast subtype composition by sample group demonstrated a marked and selective expansion of the CTHRC1+ subpopulation in both IPF upper and lower lobe samples relative to age-matched controls, whereas adventitial and alveolar subpopulations were proportionally reduced in IPF samples (**Fig. 1g**).

To validate these findings in situ and establish the spatial context of fibroblast subtype distribution, we performed multiplex spatial imaging on age-matched control and IPF lung FFPE sections (**Fig. 1h, Supplementary Fig. 1f**). In age-matched control samples (left), fibroblast-associated markers were predominantly confined to thin interstitial regions and perivascular or airway-adjacent stromal zones, with alveolar and adventitial fibroblasts constituting the dominant populations. Marker localization indicated that alveolar fibroblasts were preferentially enriched within the alveolar septa, whereas adventitial fibroblasts localized to perivascular and stromal regions, consistent with their transcriptional identities. In contrast, fibroblast foci in IPF lower lobe samples (right) displayed marked spatial disruption of normal fibroblast architecture, with enrichment of CTHRC1+ fibroblasts and relative reduction of alveolar and adventitial markers in fibrotic regions.

Spatial mapping further demonstrated that peribronchial, inflammatory, CTHRC1+, and SMC markers occupied distinct spatial domains within remodeled fibrotic tissue. CTHRC1+ and peribronchial fibroblasts appeared to have adjacent spatial localization at the borders of fibroblastic foci. Adventitial markers showed strong localization along vascular adventitia and the thickened interstitium surrounding remodeled vessels. Inflammatory fibroblast markers appeared within cell-dense stromal expansions, frequently occupying regions adjacent to inflammatory infiltrates (**Supplementary Fig. 1f**). SMCs remained spatially predominant to vascular and airway smooth muscle layers across all sample types. In particular, SMC markers exhibited a complementary spatial pattern, aligning with linear, fiber-rich structures and contractile stromal bands within fibrotic regions and were observed to border areas enriched for inflammatory fibroblasts. Together, these findings demonstrate that fibroblast heterogeneity in IPF is defined by both transcriptional reprogramming, marked by CTHRC1+ subpopulation expansion, and spatially organized architectural remodeling, where fibroblast subtypes appear to occupy distinct yet coordinated niches associated with fibrotic progression.

### Differential gene expression and fibrosis-related pathway enrichment across lung mesenchymal subtypes

To define transcriptional changes underlying fibroblast dysfunction in IPF, we performed differential gene expression (DGE) analysis comparing each fibroblast subtype from IPF upper lobe (**Supplementary Fig. 2a**) or IPF lower lobe samples (**Supplementary Fig. 2b**) to those from age-matched control samples. In IPF upper lobe samples, alveolar fibroblasts showed moderate upregulation of ECM components (*COL6A3, COL18A1, VCAN*), growth factors (*IGFBP4*), and metabolic mediators (*NNMT, SERPINF1*), with concurrent downregulation of homeostatic genes (*MACF1, TNS1, CDH13, LAMA4*) (**Supplementary Fig. 2a**). Additionally, Adventitial fibroblasts displayed constrained transcriptional responses, while inflammatory fibroblasts upregulated stromal signaling, immune/inflammatory, and matrix-remodeling-associated genes (*IGFBP4*, *COL18A1*, *SAMHD1*, *TSC22D3*, *MMP19, SFRP2, IGHG1, PTGDS*). Peribronchial fibroblasts from upper lobes exhibited a pronounced pro-fibrotic signature with upregulation of genes associated with structural collagens and matrix-remodeling components (*COL1A1, COL3A1, COL6A3, COMP, MMP2, SPARC*) and downregulation of genes associated with homeostatic regulators (*GPX3, PRELP, CCN5, PDGFRA*). CTHRC1+ fibroblasts from upper lobes displayed the most extensive reprogramming, with broad upregulation of ECM biosynthesis genes, matricellular proteins (*POSTN, COMP, CTHRC1*), and matrix-associated receptors (*MRC2, CERCAM*). In the IPF lower lobe samples, transcriptional dysregulation was substantially amplified across all subtypes, with higher fold-changes and greater numbers of DGEs. CTHRC1+ fibroblasts again showed the most profound reprogramming, with dysregulated genes including markedly elevated structural collagens and pro-fibrotic effectors (*CTHRC1, POSTN, ASPN, CTSK*), establishing this population as the most transcriptionally activated subtype in advanced IPF (**Supplementary Fig. 2b**).

Functional pathway enrichment analysis revealed consistent upregulation of ECM-related processes in fibroblast subtypes of the upper and lower lobes of IPF samples compared with age-matched control samples. Alveolar and peribronchial fibroblasts showed the most prominent enrichment of collagen biosynthesis, ECM organization, and extracellular structure organization pathways, with the highest gene ratios and statistical significance in the upper lobe samples from IPF lungs. Adventitial and inflammatory fibroblasts exhibited moderate enrichment of ECM disassembly and matrix organization terms, while SMCs showed minimal pathway activation. This pattern was largely conserved in the lower lobe, with inflammatory fibroblasts showing comparatively greater ECM-related enrichment than in the upper lobe. Notably, CTHRC1+ fibroblasts could not be included in this analysis due to an insufficient number of differentially expressed genes. These findings indicate that IPF fibroblast subtypes, particularly alveolar and peribronchial populations, adopt a pro-fibrotic, matrix-remodeling transcriptional state regardless of lobe location (**Supplementary Fig. 2c, d**).

### Collagen and matrix metalloproteinase gene expression profiles across mesenchymal subtypes in IPF

To characterize fibrotic gene expression programs, we assessed collagen family and MMP scores across fibroblast subtypes in age-matched control samples and IPF upper and lower lobe samples. Collagen family scores were significantly elevated in alveolar, inflammatory, CTHRC1+, and peribronchial fibroblasts from IPF samples relative to age-matched controls, with the most pronounced increases observed in the lower lobe (**Fig. 2a**). Adventitial fibroblasts showed a more limited increase, with significance observed only in IPF lower lobe samples, whereas SMCs showed smaller but significant increases in both IPF lobes. Heatmap analysis revealed that fibrillar, network-forming, and FACIT-associated collagens were most strongly upregulated in CTHRC1+ fibroblasts across both IPF lobes, while basement membrane-associated and anchoring collagens showed more subtype-restricted expression patterns (**Fig. 2b**).

**Figure 2.**
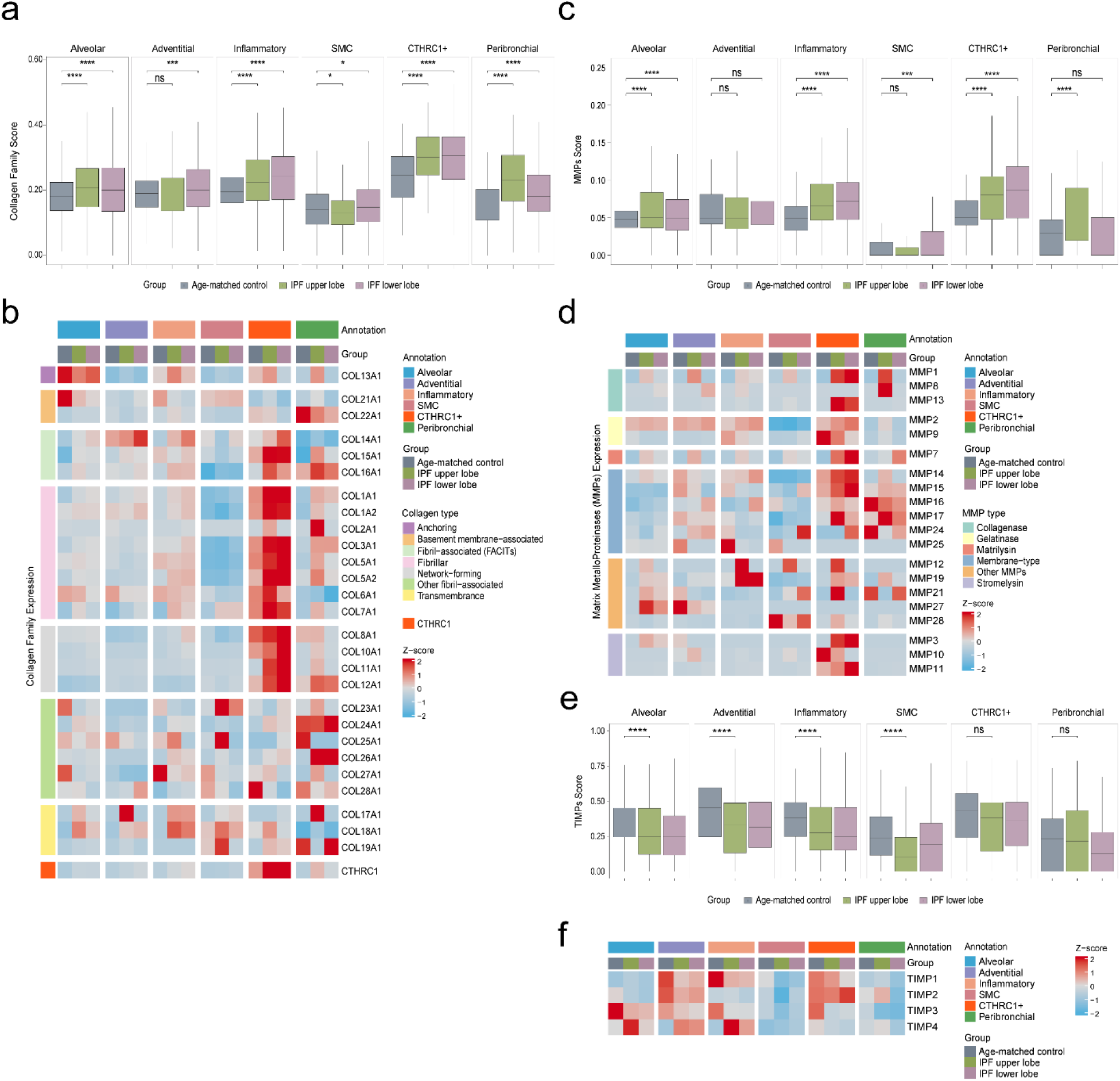
Fibrotic signatures across mesenchymal subtypes. **a.** Box plots of collagen family scores across mesenchymal subtypes and sample groups. Scores were computed using bioinformatic pipelines (see Methods). Significance is indicated as *p < 0.05, ***p < 0.001, ****p < 0.0001, and ns, not significant. **b.** Heatmap of collagen family gene expression organized by collagen category, mesenchymal subtype, and sample group. **c.** Comparative box plots illustrating MMP activity scores across mesenchymal subtypes and sample groups, including IPF explants. Scores were derived from computational analysis (see Methods). Significance is indicated as ***p < 0.001, ****p < 0.0001, and ns, not significant. **d.** Heatmap depicting expression profiles of individual MMP genes across mesenchymal subtypes and sample groups, organized by MMP class. **e.** Box plots showing TIMPs score across mesenchymal subtypes in sample groups. Statistical significance determined by Wilcoxon test; ****p < 0.0001, ***p < 0.001, ns, not significant. **f.** Heatmap showing *TIMP1*–*TIMP4* expression across mesenchymal subtypes and groups. Color scale represents Z-score normalized expression (red, high; blue, low).

MMP scores were significantly increased in alveolar, inflammatory, and CTHRC1+ fibroblasts from IPF samples compared with age-matched controls, whereas adventitial fibroblasts showed no significant differences and SMCs and peribronchial fibroblasts showed lobe-specific differences (**Fig. 2c**). At the gene level, CTHRC1+ fibroblasts showed the strongest relative expression of multiple MMP genes (*MMP1, MMP13, MMP2, MMP7, MMP14, MMP15, MMP17, MMP19, MMP3, MMP11*), with additional subtype-restricted expression of MMPs in alveolar (*MMP3, MMP19, MMP27, MMP21*), inflammatory (*MMP19, MMP14, MMP2*), and peribronchial fibroblasts (*MMP15, MMP17*) and SMCs (**Fig. 2d**). Together, these findings highlight CTHRC1+ fibroblasts as the subtype with the most prominent collagen biosynthesis and matrix remodeling transcriptional signature in IPF.

Consistent with a broadly permissive ECM remodeling environment, TIMPs score analysis revealed a significant reduction across all fibroblast subtypes in IPF upper and lower lobe relative to age-matched controls (**Fig. 2e**). Alveolar, adventitial, inflammatory, peribronchial and SMCs all showed significantly decreased TIMP scores, while CTHRC1+ fibroblasts exhibited no significant change, suggesting a subtype-specific uncoupling of protease inhibition from fibrogenic activation. At the gene level, *TIMP1* and *TIMP3* drove the bulk of this reduction, with *TIMP2* and *TIMP4* showing comparatively modest and subtype-variable changes (**Fig. 2f**). Together, these findings indicate that reduced TIMP expression across most fibroblast subtypes creates a permissive environment for MMP activity and downstream pro-fibrotic signaling in IPF lower lobe.

### Spatial collagen gene expression localizes CTHRC1+ fibroblasts to IPF fibroblastic foci and identifies them as the dominant collagen-expressing mesenchymal population

To further characterize the collagen gene expression and its distribution across fibroblast subtypes in healthy parenchyma and pathologist-annotated fibroblastic foci, we performed spatial transcriptomic analysis across age-matched control, IPF upper lobe, and IPF lower lobe samples. In age-matched control parenchyma, CTHRC1+ fibroblast markers and collagen family genes were sparsely distributed across alveoli and interstitium, consistent with a homeostatic tissue state (**Fig. 3a, left**). In contrast, fibroblastic foci in both IPF lobes showed marked spatial enrichment of CTHRC1+ fibroblast markers and collagen-associated transcripts, with dense signal concentrated within fibroblastic foci in both regions (**Fig. 3a, middle and right**).

**Figure 3.**
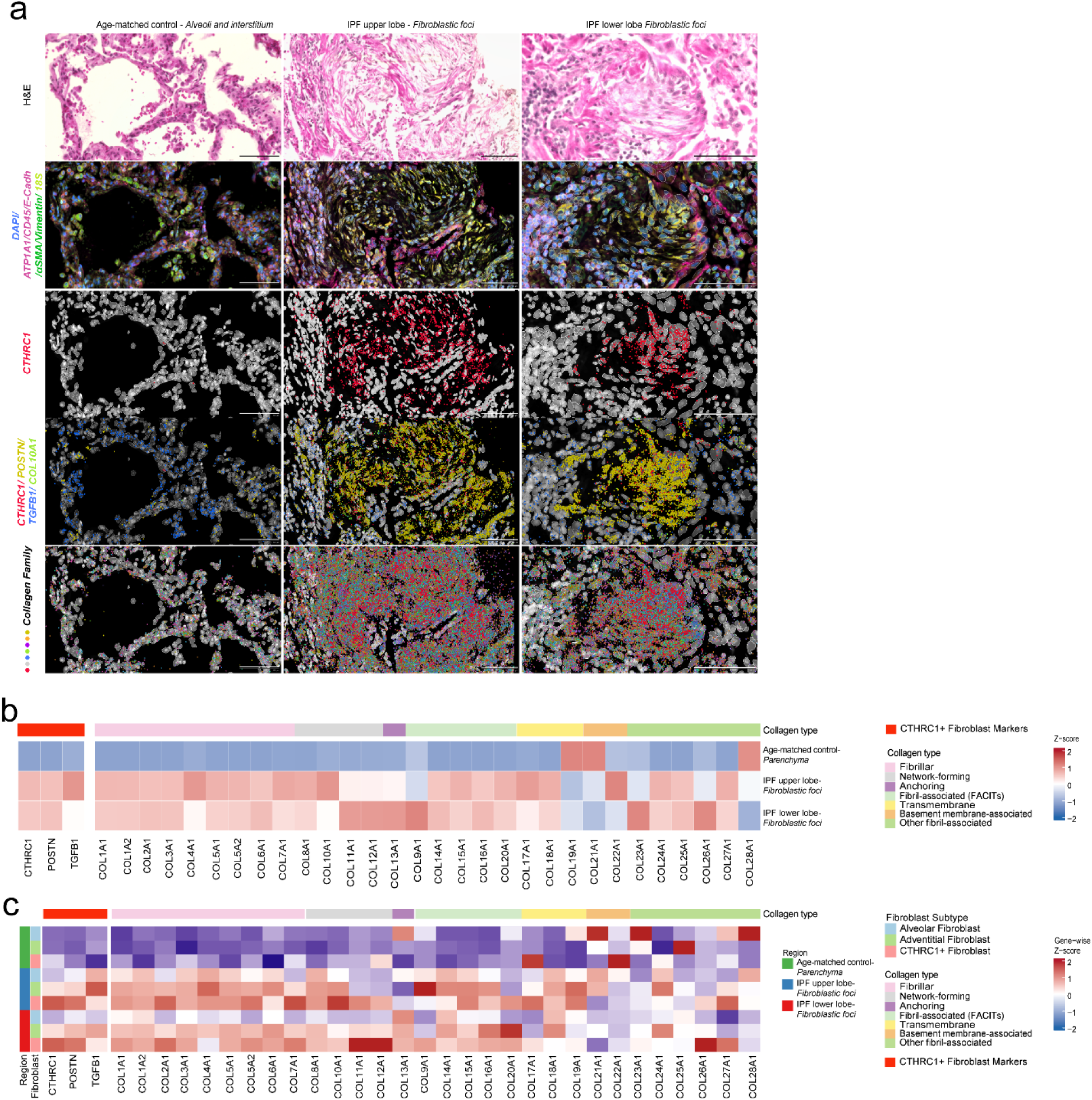
Spatial mapping and quantification of collagen family genes in mesenchymal subtypes across healthy and IPF lungs. **a.** Spatial mapping of collagen family genes and CTHRC1+ fibroblast markers in lung samples. Left, middle, and right columns show age-matched control parenchyma (alveoli and interstitium), IPF upper lobe fibroblastic foci, and IPF lower lobe fibroblastic foci, respectively. The top row shows corresponding H&E micrographs, and the remaining rows show cell segmentation and spatial localization of CTHRC1+ fibroblast markers and collagen family genes. **b.** Heat map quantifying CTHRC1+ fibroblast markers and collagen types in age-matched control parenchyma from an age-matched control sample and fibroblastic foci from IPF upper and lower lobe samples. **c.** Heat map quantifying CTHRC1+ fibroblast markers and collagen types by anatomical region and fibroblast subtypes, including alveolar, adventitial, and CTHRC1+ fibroblasts.

Heatmap quantification confirmed that CTHRC1+ fibroblast markers, fibrillar collagens (*COL1A1, COL1A2, COL3A1*), and FACIT collagens were most highly expressed within IPF fibroblastic foci compared with age-matched control parenchyma, whereas age-matched control parenchyma showed uniformly lower relative expression across most genes (**Fig. 3b**). When stratified by anatomical region and fibroblast subtype, CTHRC1+ fibroblasts within fibroblastic foci showed the highest collagen gene expression across multiple collagen classes, including fibrillar, network-forming, anchoring, FACIT, and transmembrane collagens, compared with alveolar and adventitial fibroblasts in both parenchyma and fibroblastic foci (**Fig. 3c**). Collectively, these data spatially confirm that CTHRC1+ fibroblasts are preferentially enriched within fibroblastic foci and represent the dominant fibroblast population associated with high collagen transcriptional activity in IPF lung tissue.

### CTHRC1+ and inflammatory fibroblasts emerge as central hubs of altered intercellular communication in IPF

To investigate how fibroblast subtype interactions are remodeled in IPF, we performed ligand-receptor interaction analysis across fibroblast subtypes in age-matched control, IPF upper lobe, and IPF lower lobe samples using CellChat. Global intercellular signaling analysis revealed a progressive decrease in the total number of predicted ligand-receptor interactions and overall interaction strength from age-matched control to IPF upper and lower lobes (**Fig. 4a, left**), indicating that inter-fibroblast communication networks become progressively disrupted in IPF, with the greatest impairment in the lower lobe, consistent with the known basilar predominance of fibrosis. In addition to the reduction in number of predicted interactions, the remaining signaling interactions were weaker in IPF samples (**Fig. 4a, right**). This suggests a functional loss of communication efficiency between fibroblast subtypes, reflecting fibrotic remodeling, ECM expansion, and altered cellular composition.

**Figure 4.**
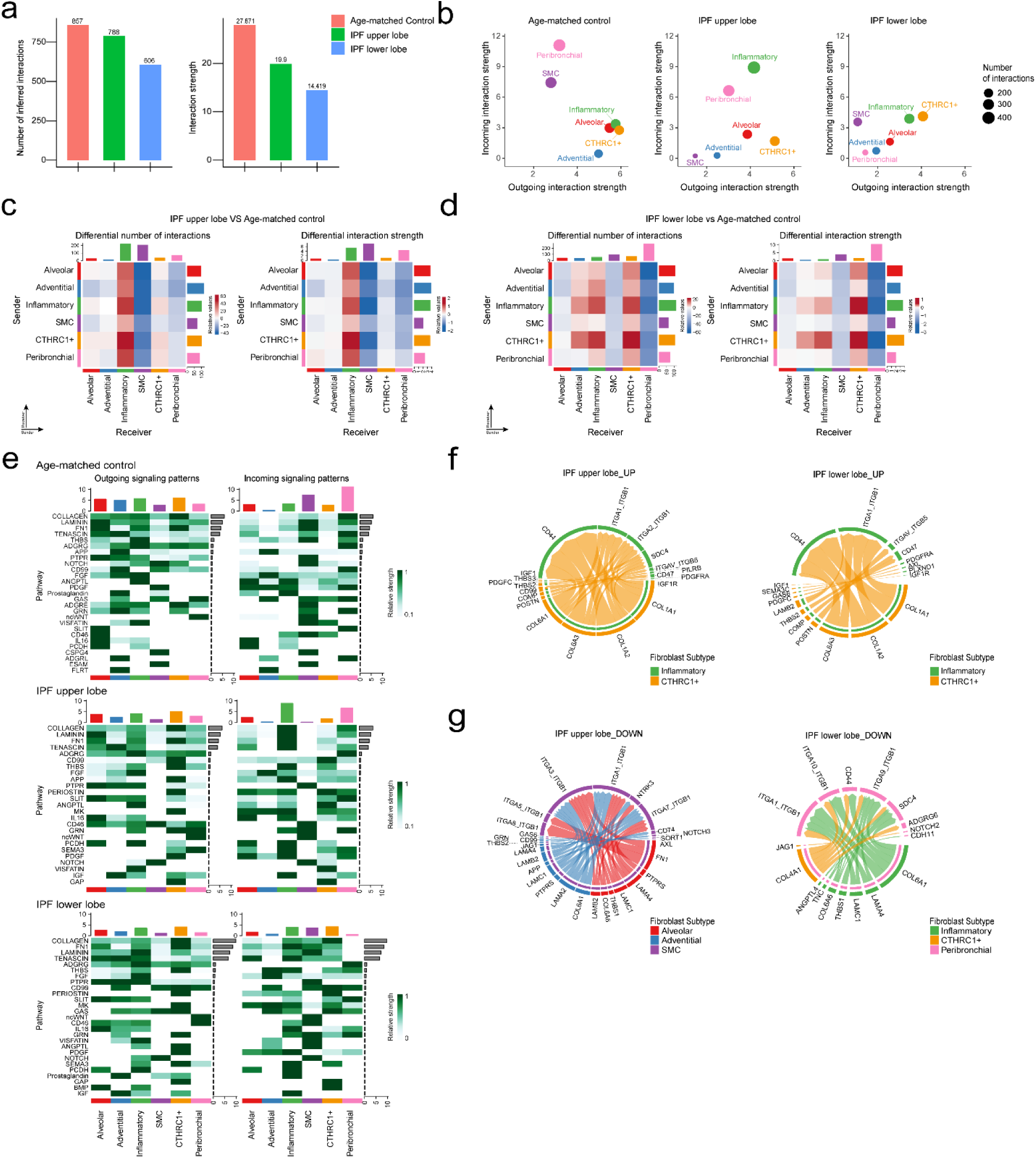
Altered cell-cell communication in IPF mesenchymal cells. **a.** Total predicted ligand-receptor interactions (left) and interaction strength (right) across age-matched control, IPF upper lobe, and IPF lower lobe samples. **b.** Scatter plots showing outgoing (x-axis) and incoming (y-axis) interaction strength among fibroblast subtypes in age-matched control (left), IPF upper lobe (middle), and IPF lower lobe (right) samples. Point size denotes number of interactions. **c.** Heatmaps showing differential number of interactions (left) and differential interaction strength (right) between mesenchymal subtypes in IPF upper lobe samples compared with age-matched control samples. Receiver mesenchymal subtypes are displayed on the x-axis and sender mesenchymal subtypes are displayed on the y-axis. Color intensity represents effect size. **d.** Corresponding differential interaction patterns between mesenchymal subtypes in IPF lower lobe versus age-matched control. Receiver mesenchymal subtypes are displayed on the x-axis and sender mesenchymal subtypes are displayed on the y-axis. Color intensity represents effect size. **e.** Pathway-level signaling patterns. Outgoing and incoming signaling activities are shown side-by-side for age-matched control (top), IPF upper lobe (middle), and IPF lower lobe (bottom). Columns correspond to mesenchymal subtypes. For each panel, the colored bars above the heatmap indicate the summed outgoing or incoming signaling strength for that mesenchymal population, while the bars to the right summarize total pathway activity across all mesenchymal subtypes. Rows represent individual signaling pathways. Color scale reflects relative signaling strength through row-scaled. **f.** Circular chord diagrams depicting upregulated ligand-receptor interactions among mesenchymal subtypes in IPF upper lobe (left) and IPF lower lobe (right) samples compared with age-matched control samples. Each arc along the outer ring represents an individual ligand or receptor gene, and chords denote directional ligand-receptor connections. Colors indicate the mesenchymal subtype contributing to each interaction. **g.** Circular chord diagrams depicting downregulated ligand-receptor interactions among mesenchymal subtypes in IPF upper lobe (left) and IPF lower lobe (right) samples compared with age-matched control samples. Each arc along the outer ring represents an individual ligand or receptor gene, and chords denote directional ligand-receptor connections. Colors indicate the mesenchymal subtype contributing to each interaction.

Scatter plot analysis of outgoing versus incoming interaction strength showed that CTHRC1+ fibroblasts exhibited the highest outgoing interaction strength in both IPF lobes (**Fig. 4b**). In IPF upper lobe samples, inflammatory fibroblasts exhibited the highest incoming interaction strength, whereas in IPF lower lobe samples, CTHRC1+ fibroblasts showed slightly higher incoming interaction strength than inflammatory fibroblasts. By contrast, interaction patterns in age-matched control samples were more broadly distributed across subtypes. Although alveolar fibroblasts are widely accepted as precursors of multiple fibroblast lineages, this lineage relationship did not correspond to dominant communication hub status in the steady-state, non-injured lung.^11^ In age-matched control samples, the network analysis showed that peribronchial fibroblasts and SMCs both sent and received ligand-receptor traffic, but their roles were more prominent as receivers, whereas inflammatory, alveolar, and CTHRC1+ fibroblasts exhibited relatively stronger outgoing interaction strength. This hub-like organization reflects active signaling demand, tissue niches, and homeostatic roles. In contrast, IPF samples showed a marked redistribution of communication roles. Consistent with their role as key mediators of fibrotic ECM remodeling,^12^ CTHRC1+ fibroblasts emerged as dominant signaling hubs with the highest outgoing interaction strength in both lobes and high incoming interaction strength in the lower lobe. Inflammatory fibroblasts emerged as the dominant incoming state in the IPF upper lobe samples and remained a prominent incoming state in the lower lobe samples. Inflammatory fibroblasts became prominent signaling sources and targets, reflecting persistent injury responses, immune activation, and cytokine-driven crosstalk previously described in IPF pathogenesis. Peribronchial and SMCs lost their dominant communication roles in IPF, consistent with the destruction and remodeling of normal lung architecture.^13^ Persistent outgoing signaling from alveolar fibroblasts further suggested chronic cellular stress and dysfunctional epithelial-mesenchymal crosstalk.^14,15^ Collectively, these findings indicate that IPF shifts fibroblast communication from a broadly distributed homeostatic organization toward a more polarized pro-fibrotic and inflammatory network dominated by CTHRC1+ and inflammatory fibroblast populations.

Differential interaction analysis comparing age-matched control samples with IPF upper lobe (**Fig. 4c**) and IPF lower lobe (**Fig. 4d**) samples revealed that CTHRC1+ fibroblasts emerged as the primary gainers of outgoing interactions in IPF upper lobe samples, with the strongest gains directed toward inflammatory receiver states compared with age-matched control. Inflammatory fibroblasts gained the most incoming interactions, becoming a dominant signaling target. In contrast, interactions directed toward SMCs were broadly reduced, indicating a loss of communication prominence, with the most consistent reductions observed for SMC receiver states (**Fig. 4c, left**). Differential interaction strength analysis largely mirrored these patterns, with CTHRC1+ fibroblasts showing the strongest gain in outgoing signaling strength directed toward adventitial and inflammatory subtypes, while peribronchial and SMC receiver states exhibited reduced interaction strength across most sender-receiver pairs (**Fig. 4c, right**).

Communication reorganization was most pronounced in IPF lower lobe samples, where CTHRC1+ fibroblasts showed the greatest gains in outgoing interaction number and strength, broadly targeting adventitial, inflammatory, and CTHRC1+ receiver states (**Fig. 4d**). Inflammatory fibroblasts remained prominent signaling recipients, while peribronchial fibroblasts and SMCs consistently lost communication prominence. Collectively, these data reveal a lobe-dependent and progressive rewiring of mesenchymal signaling networks in IPF, driven by CTHRC1+ sender dominance and inflammatory fibroblast centrality.

To characterize pathway-level signaling changes across fibroblast subtypes, we analyzed outgoing and incoming signaling patterns in age-matched control, IPF upper lobe, and IPF lower lobe (**Fig. 4e**). Pathway activity was redistributed across groups, with age-matched control showing broader distribution across fibroblast subtypes and IPF upper and lower lobe samples showing greater concentration of signaling activity within specific fibroblast populations. In age-matched control samples, outgoing signaling was broadly distributed across fibroblast subtypes with collagen, laminin, tenascin, and fibronectin pathways showing relatively uniform activity across alveolar, adventitial, and peribronchial fibroblasts and SMCs. Incoming signaling patterns were similarly distributed, with stronger incoming activity in SMC and peribronchial fibroblasts but no single subtype dominating the communication landscape, consistent with homeostatic tissue maintenance (**Fig. 4e, top**). In IPF upper lobe samples, outgoing signaling patterns were markedly reorganized. CTHRC1+ fibroblasts emerged as the dominant senders of collagen, fibronectin, periostin, THBS, and PDGF pathways relative to age-matched control. Alveolar and inflammatory fibroblasts showed increased outgoing signaling through inflammation-associated pathways (GRN, IL16, ANGPTL, IGF). Incoming signaling patterns revealed inflammatory and peribronchial fibroblasts as the primary recipients of amplified signals, while adventitial and SMCs showed relatively reduced incoming pathway activity (**Fig. 4e, middl**e). In IPF lower lobe samples, these patterns were further amplified, with CTHRC1+ and inflammatory fibroblasts displaying the strongest outgoing signaling across collagen, periostin, fibronectin, laminin, and additional remodeling pathways (TENASCIN, THBS, NOTCH, PDGF, SLIT). Incoming signaling to CTHRC1+, inflammatory, and SMCs showed greater prominence in the lower lobe, reinforcing their roles as central communication hubs in disease (**Fig. 4e, bottom**). Collectively, these data demonstrate a progressive, lobe-dependent remodeling of pathway-level signaling in IPF, with CTHRC1+ fibroblasts acquiring dominant outgoing signaling roles and inflammatory fibroblasts becoming primary signaling recipients, fundamentally disrupting the balanced communication network observed in age-matched control samples.

Beyond cell-cell interactions, we also resolved ECM-specific communication probabilities for collagen, laminin, FN1, and tenascin pathways across sender-receiver pairs in fibroblast subtypes (**Supplementary Fig. 3**). Analysis of predicted collagen ligand-receptor communication probabilities across fibroblast subtypes revealed a progressive redistribution and concentration toward CTHRC1+ fibroblasts in IPF (**Supplementary Fig. 3a**). In IPF samples, collagen signaling became increasingly concentrated, with CTHRC1+ fibroblasts emerging as the dominant sender population compared with age-matched control samples, where the collagen-mediated communication was broadly distributed across multiple sender-receiver pairs. In IPF upper lobe samples, CTHRC1+ fibroblasts directed amplified collagen communication toward inflammatory, peribronchial, and alveolar receiver subtypes, and in the IPF lower lobe, this redistribution remained evident across multiple receiver subtypes. These findings indicate that IPF disrupts the balanced collagen signaling network observed in homeostatic lung tissue, driving a pathological concentration of collagen-mediated intercellular communication through CTHRC1+ fibroblasts in a lobe-dependent manner. Laminin ligand-receptor communication probabilities revealed that, in the IPF upper lobe, there was a marked reduction across most sender-receiver pairs, with selective retention in alveolar and CTHRC1+ fibroblasts. This reduction was further pronounced in the IPF lower lobe, where laminin communication probabilities were broadly diminished across multiple fibroblast subtype interactions, suggesting progressive loss of basement membrane-associated homeostatic signaling with advancing fibrotic remodeling compared with age-matched control samples (**Supplementary Fig. 3b**). These findings indicate that IPF is associated with a progressive deterioration of laminin-mediated intercellular communication, reflecting the disruption of normal basement membrane architecture characteristic of advanced fibrotic lung disease. Analysis of FN1 ligand-receptor communication probabilities revealed progressive amplification and redistribution of this signaling in IPF (**Supplementary Fig. 3c**), reflecting its role in cell adhesion, matrix assembly, and tissue repair. In IPF upper lobe samples, FN1 signaling was redistributed toward CTHRC1+, alveolar, and inflammatory fibroblasts as dominant senders, with inflammatory autocrine signaling and alveolar fibroblasts emerging as prominent receivers.

Consistently, in the lower lobe, CTHRC1+ fibroblasts displayed the highest FN1 outgoing signaling strength. Tenascin, a matricellular glycoprotein that regulates fibroblast migration signaling,^16^ showed amplified communication probabilities in IPF samples, whereas in age-matched control samples it showed modest activity concentrated in alveolar and adventitial senders and in peribronchial, SMC, and alveolar receivers, consistent with its homeostatic role in tissue elasticity and matrix organization. In IPF upper lobe samples, tenascin signaling became selectively upregulated in alveolar, adventitial, and inflammatory fibroblasts, with increased probabilities directed toward alveolar and inflammatory receivers, establishing an autocrine signaling loop. Consistent with these trends, in the lower lobe tenascin communication probabilities reached their highest levels, with inflammatory fibroblasts displaying broader and stronger outgoing signaling across multiple receiver subtypes than in the upper lobe, except toward adventitial fibroblasts (**Supplementary Fig. 3d**). In all, these ECM-focused analyses extend the pathway-level changes summarized in **Fig. 4e** and are consistent with the ECM– and integrin-centered rewiring highlighted in Fig. 4f, g, indicating that progressive amplification of collagen and FN1 signaling centered on CTHRC1+ fibroblasts, together with increased tenascin signaling and loss of laminin-mediated homeostatic communication, reflecting a fundamental and lobe-dependent disruption of ECM signaling balance underlying pathological fibrotic remodeling in IPF.

Finally, to identify specific ligand-receptor pairs underlying the reorganization of intercellular communication in IPF, we examined upregulated and downregulated interactions across fibroblast subtypes in IPF upper and lower lobe samples compared with age-matched control samples (**Fig. 4f, g**). In both IPF upper and lower lobe samples, upregulated ligand-receptor interactions were predominantly driven by CTHRC1+ fibroblasts, which contributed the largest proportion of gained interactions (Fig. 4f). The most prominent upregulated pairs involved collagen-integrin signaling axes (*COL1A1, COL1A2, COL6A3, COL6A1* interacting with *ITGA1_ITGB1, ITGA2_ITGB1, ITGAV_ITGB5*), alongside matricellular mediators (*POSTN, THBS2, COMP*) and growth factor-receptor pairs (*IGF1R, PDGFRA, SDC4*). Inflammatory fibroblasts contributed to a secondary set of upregulated interactions, particularly involving *CD44* and integrin-associated ligands. The lower lobe showed a broader repertoire of upregulated interactions compared with the upper lobe, with additional pairs involving *SEMA3A, LAMB2,* and *PDGFC*, consistent with more advanced fibrotic remodeling. Downregulated interactions in IPF upper lobe samples were distributed across alveolar, adventitial, and SMCs, involving laminin-integrin pairs (*LAMB2, LAMC1, LAMA2, LAMA4* with *ITGA3_ITGB1, ITGA1_ITGB1, ITGA7_ITGB1*), homeostatic signaling mediators (*FN1, APP, GRN*), and cell-cell communication receptors (*NOTCH3, SORT1, AXL, CD74*) (**Fig. 4g**). In IPF lower lobe samples, downregulated interactions were primarily attributed to inflammatory, CTHRC1+, and peribronchial fibroblasts, with loss of collagen-integrin (*COL4A1*) and Notch signaling pairs (*NOTCH2, JAG1*) (**Fig. 4g**). Collectively, these data demonstrate that IPF is characterized by a fundamental rewiring of ligand-receptor communication, with CTHRC1+ fibroblasts acquiring dominant roles in collagen-integrin and matricellular signaling, while homeostatic laminin, fibronectin, and Notch-mediated interactions are progressively lost across multiple fibroblast subtypes in a lobe-dependent manner.

### Pseudotime trajectory analysis reveals progressive transcriptional reprogramming toward a CTHRC1+ pro-fibrotic state in IPF

To investigate the transcriptional dynamics underlying fibroblast state transitions in IPF, we performed pseudotime trajectory and partition-based graph abstraction (PAGA) analyses across fibroblast subtypes in age-matched control, IPF upper lobes, and IPF lower lobes. UMAP embedding revealed spatially distinct but partially continuous fibroblast subtype clusters suitable for trajectory inference (**Fig. 5a**). PAGA analysis revealed a structured fibroblast connectivity topology in which CTHRC1+ fibroblasts showed the strongest transcriptional connectivity with inflammatory and peribronchial fibroblasts, suggesting shared transcriptional programs or close relationships between these populations (**Fig. 5b**). Inflammatory fibroblasts showed intermediate connectivity bridging CTHRC1+ and alveolar fibroblasts, while adventitial fibroblasts and SMCs occupied peripheral positions with weaker network connectivity, consistent with their transcriptionally distinct identities.

**Figure 5.**
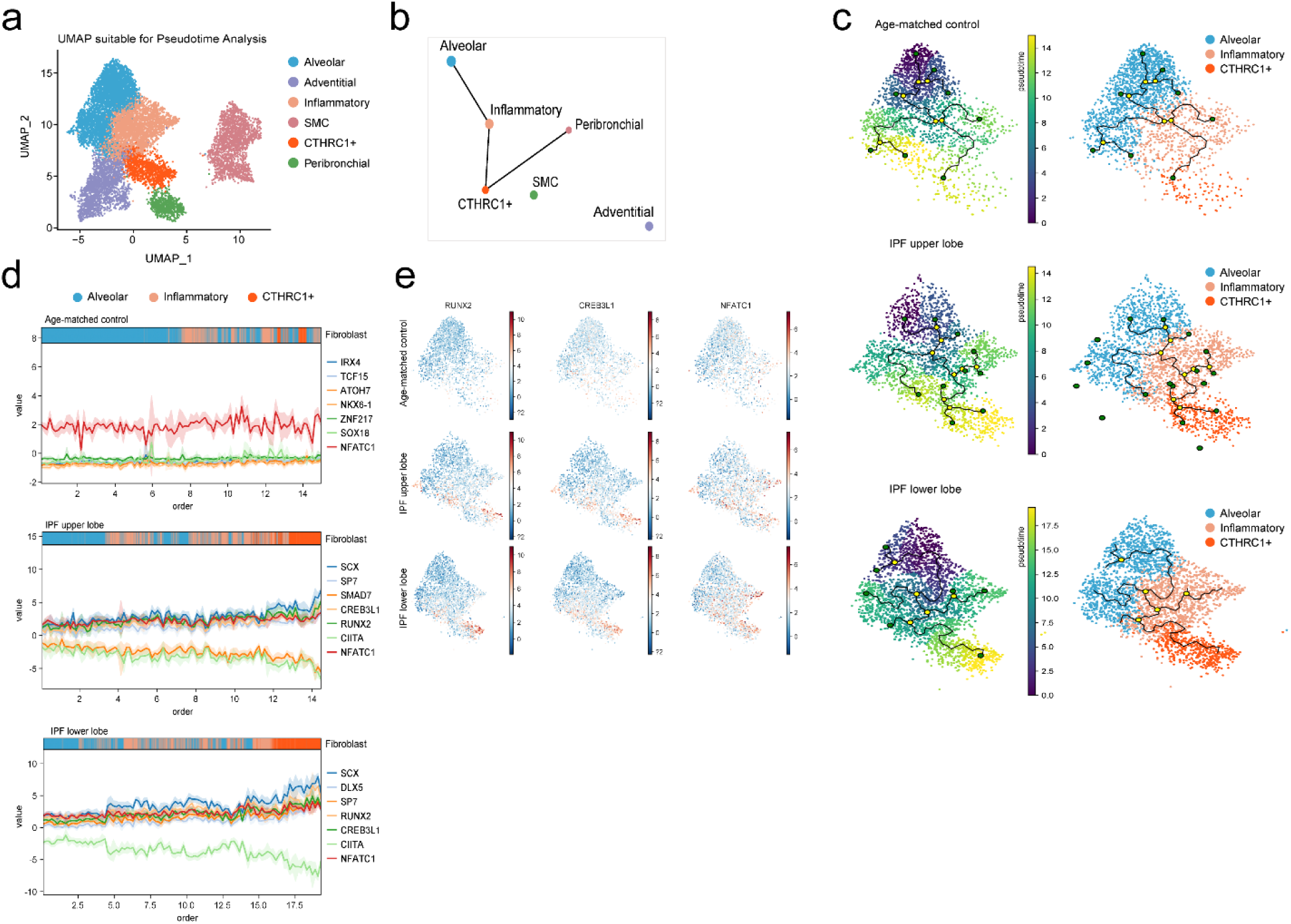
Connectivity and transcriptional trajectories of mesenchymal subtypes in IPF lung tissue. **a.** UMAP embedding of lung tissue-derived mesenchymal subtypes colored by subtype identity: alveolar (blue), adventitial (light blue), inflammatory (orange), SMC (peach), CTHRC1+ (red), and peribronchial (green). Spatial relationships between clusters informed downstream pseudotime trajectory inference. **b.** PAGA graph showing inferred connectivity among mesenchymal subtypes, highlighting inferred relationships among mesenchymal subtypes. **c.** Monocle trajectory analysis showing transcriptional progression of mesenchymal subtypes with alveolar fibroblasts positioned at the transcriptional root of the inferred trajectory, across age-matched control, IPF upper lobe, and IPF lower lobe samples. Left panel: Cells ordered along pseudotime, colored by progression from early (yellow) to late (dark blue) states; black lines represent the trajectory backbone, and yellow nodes indicate branch points corresponding to transcriptional divergence. Right panel: Cells colored by cluster identity, including alveolar, inflammatory and CTHRC1⁺ fibroblasts. Trajectories reveal potential differentiation paths and lineage relationships among three fibroblast subtypes in IPF compared to age-matched control lung samples. **d.** TF activity dynamics along pseudotime trajectory using the CollecTRI database in age-matched control, IPF upper and lower lobes. Lines represent smoothed TF activity values with shaded confidence intervals; color bars indicate fibroblast subtype composition along the trajectory. Representative TFs for each group are shown at right. **e.** UMAP visualization of inferred TF activity scores for RUNX2, CREB3L1, and NFATC1 across the combined dataset and within age-matched control, IPF upper, and IPF lower lobe samples. Rows correspond to group; columns correspond to TF. Color scale indicates TF activity from low (blue) to high (red).

Based on the strong transcriptional connectivity, shared pro-fibrotic gene expression programs, and dominant intercellular communication interactions identified among alveolar, inflammatory, and CTHRC1+ fibroblasts, we performed pseudotime trajectory inference and transcription factor (TF) activity analysis along the inferred trajectory across age-matched control, IPF upper lobe, and IPF lower lobe samples to further resolve the transcriptional dynamics underlying fibroblast state transitions in IPF (**Fig. 5c**). In age-matched control samples, cells were broadly distributed across early pseudotime, with CTHRC1+ fibroblasts representing a minor terminal population. In IPF upper lobe samples, an increased proportion of cells advanced toward later pseudotime states enriched in CTHRC1+ fibroblasts. This pattern was most pronounced in the IPF lower lobe, where the majority of cells accumulated at terminal pseudotime positions dominated by CTHRC1+ fibroblasts, consistent with progressive fibroblast reprogramming in advanced disease.

TF activity analysis along the pseudotime trajectory using the CollecTRI database identified group-specific transcriptional regulators associated with driving fibroblast state transitions (**Fig. 5d**). In age-matched control, homeostatic TFs including IRX4, TCF15, and NKX6-1 showed relatively stable activity. In IPF upper lobe, pro-fibrotic TFs including SCX, RUNX2, SP7, and CREB3L1 emerged with increasing activity along pseudotime, while NFATC1 showed progressive increases in activity across all groups. In the lower lobe, SCX, RUNX2, and DLX5 displayed the strongest pseudotime-dependent increases in activity, with CIITA showing consistent downregulation, suggesting suppression of immune-regulatory transcriptional programs in advanced fibrosis. UMAP visualization of TF activity confirmed that RUNX2, CREB3L1, and NFATC1 activity was progressively enriched in IPF compared to age-matched control, with the highest scores concentrated in CTHRC1+ fibroblasts in the lower lobe (**Fig. 5e**). Collectively, these findings demonstrate that IPF fibroblasts undergo directional and progressive transcriptional reprogramming toward a CTHRC1+-dominated pro-fibrotic terminal state, driven by coordinated activation of ECM-regulatory transcription factors including RUNX2, CREB3L1, SCX, and NFATC1, with the most advanced reprogramming occurring in the lower lobe, consistent with greater disease severity in this region.

### Mesenchymal subtype composition and collagen expression profiles across isolation protocols in IPF and age-matched control lung tissue

To determine whether the mesenchymal transcriptional heterogeneity and pro-fibrotic signatures identified in IPF lung samples could be recapitulated and further investigated in vitro, we characterized mesenchymal subtype composition and collagen expression profiles in cultured human primary lung fibroblasts derived from fresh tissue from the same cohorts, including age-matched control, IPF upper lobe, and IPF lower lobe samples, using three isolation protocols, whole lung cell suspension (WLCS), negative fraction, and outgrowth (**Fig. 6a**). To assess the influence of isolation methodology on subtype representation and transcriptional profiles, principal component analysis (PCA) revealed protocol-dependent transcriptional variation across all sample groups, with initial (P1) and final passage (P6) cells clustering distinctly within each protocol and condition (**Fig. 6b**). WLCS and negative fraction protocols yielded broader transcriptional dispersion compared to outgrowth, suggesting greater cellular heterogeneity captured by suspension-based approaches. In addition, IPF-derived fibroblasts showed markedly greater transcriptional dispersion between initial and final passages across WLCS and negative fraction protocols, suggesting that serial passaging induces more pronounced phenotypic drift in IPF fibroblasts. Unsupervised clustering of cultured fibroblasts identified seven transcriptionally distinct subtypes: alveolar 1, alveolar 2, adventitial, inflammatory, SMC, senescence-like, and proliferating subtypes (**Fig. 6c**), each validated by subtype-specific marker gene expression encompassing collagen family genes, matrix remodeling enzymes, and senescence and proliferation markers (**Fig. 6d**). Integrated UMAP analysis across all sample groups, isolation protocols, and passages showed broad intermixing of cells from multiple samples, without complete segregation by sample group (**Supplementary Fig. 4a**). Protocol-stratified analysis confirmed that all seven fibroblast subtypes were captured across WLCS, negative fraction, and outgrowth protocols in the available protocol and sample-group combinations, although outgrowth cultures were not available for IPF lower lobe samples (**Supplementary Fig. 4b**). Validation of cultured fibroblast subtype identity using tissue-derived mesenchymal marker genes revealed partial transcriptional correspondence between in vitro and in vivo signatures; however, a subset of tissue-specific markers showed reduced expression in cultured fibroblasts (**Supplementary Fig. 4c**), suggesting that standard in vitro culture conditions only partially preserve the transcriptional identity of lung fibroblast subtypes as defined in native tissue. Notably, two transcriptionally distinct alveolar populations were identified, alveolar 1 and alveolar 2, where alveolar 2 displayed an activated transcriptional state characterized by marked upregulation of collagens (*COL1A1, COL1A2, COL3A1, COL5A1*), consistent with a pro-fibrotic activated alveolar fibroblast phenotype.

**Figure 6.**
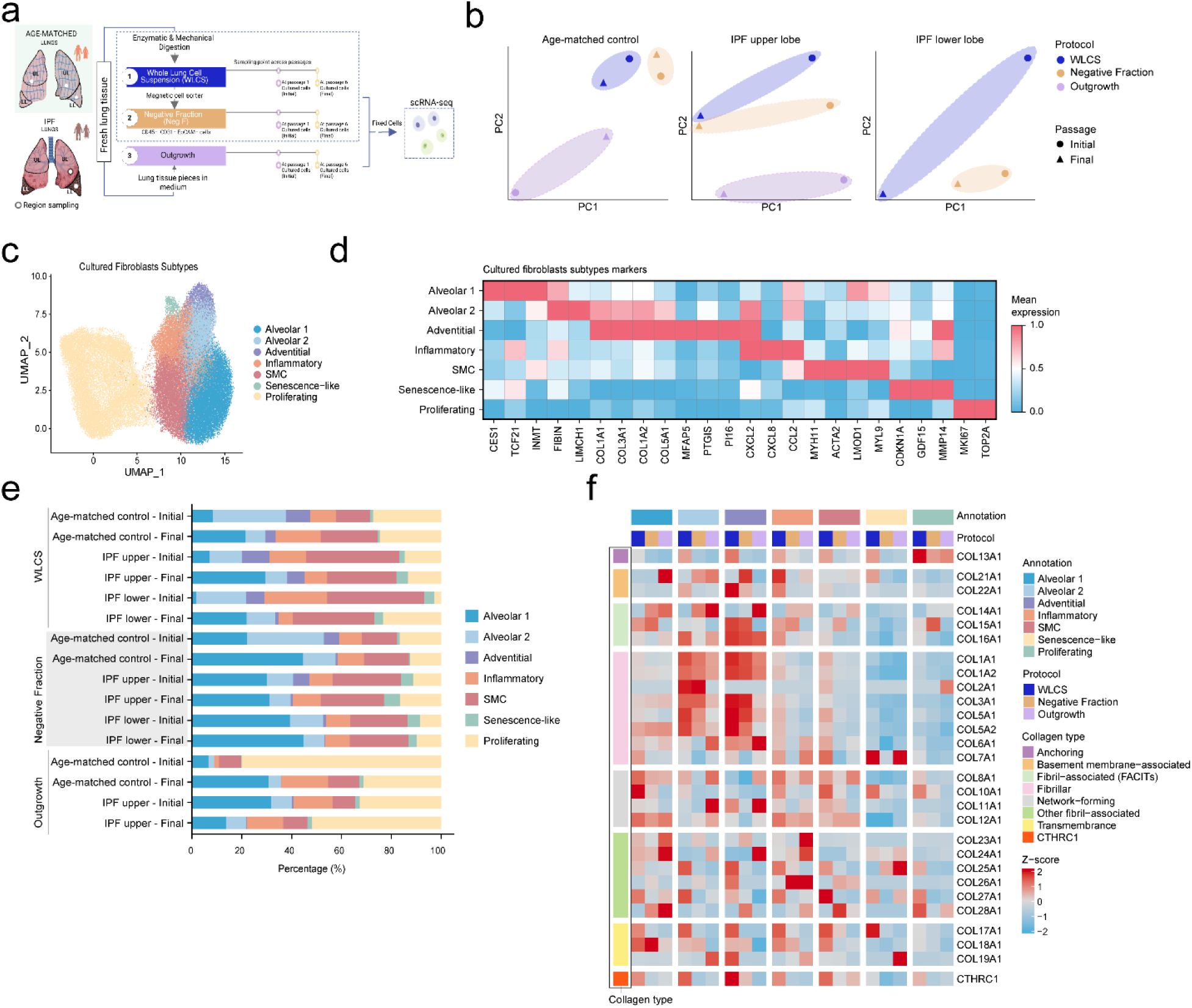
Impact of isolation protocol and in vitro culture on mesenchymal heterogeneity. **a.** Experimental design for mesenchymal isolation using three protocols: WLCS, negative fraction, and outgrowth. **b.** PCA plots comparing fibroblast samples under different conditions, isolation protocols, and passages. Colors denote isolation protocol, and symbols denote passage. **c.** UMAP visualization of fibroblast subtypes in cultured cells. **d.** Heatmap of marker gene expression across cultured fibroblast subtypes. **e.** Stacked bar plots showing the composition of fibroblast subtypes in culture across isolation protocols, sample groups, and passages. **f.** Heatmap of collagen family gene expression by collagen type, fibroblast subtype, and isolation protocol, and sample group.

Fibroblast subtype composition varied substantially across isolation protocols and conditions (**Fig. 6e**). WLCS and negative fraction protocols enriched for alveolar, inflammatory and SMC subtypes, while outgrowth cultures were predominantly composed of alveolar and proliferating fibroblasts. IPF samples showed increased proportions of senescence-like fibroblasts compared to age-matched control samples across all protocols, consistent with the known role of cellular senescence in IPF pathogenesis.^17,18^ CTHRC1+ fibroblasts were notably absent or markedly underrepresented across all cultured conditions and protocols, suggesting selective loss of this subtype under standard in vitro culture conditions.

Collagen family expression analysis across isolation protocols and conditions revealed both protocol-dependent and disease-dependent transcriptional variation (**Fig. 6f**). Basement membrane-associated, FACIT, and fibrillar collagens were the most highly expressed collagen types across all protocols, with the highest expression levels concentrated in adventitial and activated alveolar (alveolar 2) fibroblasts, consistent with their transcriptionally activated pro-fibrotic state. Network-forming and other fibril-associated collagens showed intermediate expression levels predominantly in alveolar 1, inflammatory, and SMCs, suggesting a more moderate matrix remodeling program in these subtypes. Anchoring and transmembrane collagens displayed the most variable expression patterns, showing greater sensitivity to isolation protocol and donor condition, which may reflect their context-dependent roles in basement membrane maintenance and cell-matrix interactions. Notably, *CTHRC1* expression was detected at low levels across cultured conditions, further supporting the preferential loss of CTHRC1+ fibroblasts during in vitro culture. Collectively, these findings demonstrate that fibroblast subtype composition and collagen transcriptional programs in cultured lung fibroblasts are influenced by both isolation protocol and disease state, and highlight the critical limitation that CTHRC1+ fibroblasts, identified as the most transcriptionally activated pro-fibrotic subtype in IPF tissue, are poorly represented in standard in vitro culture systems, underscoring the importance of tissue-based single-cell approaches for capturing the full spectrum of fibroblast heterogeneity in IPF.

To further resolve sample group-, isolation protocol– and passage-dependent collagen family expression patterns across cultured fibroblast subtypes, we examined collagen gene expression separately for each isolation protocol (**Supplementary Fig. 5**). In WLCS and negative fraction protocols, collagen family expression was often higher at initial than at final passage across multiple fibroblast subtypes, particularly for anchoring, FACIT, fibrillar, and network-forming collagens across multiple cultured fibroblast states (**Supplementary Fig. 5a, b**). Across both WLCS and negative fraction protocols, alveolar 2 and adventitial fibroblasts showed consistently strong expression of multiple fibrillar and FACIT-associated collagens across passages. Alveolar 2 fibroblasts showed relatively higher of fibrillar collagen expression than alveolar 1 across passages, consistent with a more activated matrix-producing state, suggesting selective activation of this subtype during culture expansion. Adventitial fibroblasts consistently maintained high FACIT and fibrillar collagen expression across both passages, reflecting a stable pro-fibrotic transcriptional identity in vitro, while alveolar 1, inflammatory, and SMCs retained network-forming collagen expression across passages. Proliferating fibroblasts isolated by negative fraction protocol showed increased expression of *COL13A1* and selected basement membrane-associated collagens, while other fibril-associated and transmembrane collagens displayed greater inter-passage variability. Notably, CTHRC1 expression was generally reduced during culture across fibroblast subtypes, although adventitial fibroblasts retained comparatively higher CTHRC1 expression, particularly in IPF-derived samples, suggesting a subtype-specific capacity to partially preserve this pro-fibrotic marker under standard culture conditions. In the outgrowth protocol, collagen family expression patterns were broadly consistent with those observed in WLCS and negative fraction protocols, but with more uniform distribution across fibroblast subtypes and smaller inter-passage differences, suggesting that outgrowth conditions partially attenuate subtype-specific collagen transcriptional programs (**Supplementary Fig. 5c**). IPF upper lobe-derived fibroblasts maintained higher fibrillar collagen expression compared to age-matched control across both passages. Notably, while *CTHRC1* expression remained low during culture in age-matched control-derived fibroblasts, IPF upper lobe samples showed a progressive relative enrichment of *CTHRC1* expression across passages, representing a unique feature of the outgrowth protocol not observed in WLCS or negative fraction conditions. However, expression remained low in outgrowth cultures overall, despite detectable expression in selected IPF upper lobe states.

To complement the collagen family expression analysis and provide a comprehensive view of ECM remodeling programs in cultured fibroblasts, we next examined MMP expression patterns, the primary enzymatic mediators of collagen degradation and matrix turnover, across isolation protocols, conditions, and passages (**Supplementary Fig. 6**). MMP expression analysis across isolation protocols revealed both shared and protocol-specific patterns of culture-associated increases. Across all three protocols, membrane-type metalloproteinases were consistently upregulated during culture in alveolar 1, alveolar 2, and adventitial fibroblasts, and stromelysin expression was observed in alveolar 1, inflammatory, and senescence-like fibroblasts, representing the most reproducible MMP responses independent of isolation methodology. Protocol-specific differences were also evident. Gelatinase expression was most broadly distributed in the WLCS protocol, affecting alveolar 1, alveolar 2, adventitial, and inflammatory fibroblasts, while in the negative fraction protocol it was restricted to adventitial and inflammatory subtypes, and was less prominent in the outgrowth protocol. Collagenase expression was most evident in senescence-like and proliferating fibroblasts in the negative fraction protocol, while matrilysin expression in inflammatory fibroblasts was shared between negative fraction and outgrowth protocols but absent in WLCS. Notably, the outgrowth protocol induced the broadest membrane-type metalloproteinase expression pattern, extending across all seven fibroblast subtypes, a pattern not observed in either WLCS or negative fraction protocols. Collectively, these findings indicate that while core MMP programs are consistently acquired during culture expansion across protocols, isolation methodology substantially influences the breadth and subtype specificity of MMP expression, with the outgrowth protocol driving the most extensive matrix remodeling transcriptional response.

### Cultured fibroblast subtypes display subtype-specific transcriptional programs enriched for ECM remodeling, inflammatory, and proliferative biological processes

To further characterize the transcriptional identity of cultured fibroblast subtypes identified in the integrated dataset, we performed differential gene expression (DGE) analysis and Gene Ontology (GO) biological process enrichment for each subtype (**Supplementary Fig. 7**), complementing the subtype marker, collagen, and MMP signatures presented in **Fig. 6** and **Supplementary Figs. 5-6**.

Alveolar 1 and alveolar 2 fibroblast subtypes were both enriched for ECM organization; alveolar 1 fibroblast were additionally enriched for extracellular structure organization and regulation of cell proliferation and migration, consistent with a stromal-mesenchymal transcriptional identity, whereas alveolar 2 fibroblasts showed stronger enrichment for collagen fibril organization (**Supplementary Fig. 7a, b**). Alveolar 2 fibroblasts also displayed a more pronounced pro-fibrotic gene expression profile, with preferential upregulation of fibrillar collagens (*COL1A1, COL1A2, COL3A1, COL5A1*), TGFβ-associated matricellular genes (*THBS2, SPARC*) and the transcription factor *GATA6*, distinguishing them from the alveolar 1 subtype and suggesting a transcriptional state associated with active ECM remodeling (**Supplementary Fig. 7a, b**). Adventitial fibroblasts were enriched for collagen fibril organization, extracellular matrix structure organization, and regulation of cell population proliferation, reflecting their known roles in perivascular connective tissue maintenance (**Supplementary Fig. 7c**). Inflammatory fibroblasts showed prominent enrichment for biological processes, including cytokine-mediated signaling, cellular response to cytokine stimulus, and negative regulation of apoptotic processes (**Supplementary Fig. 7d**), consistent with an activated stromal inflammatory phenotype. SMC displayed the most distinct transcriptional profile, with strong enrichment for collagen fibril organization, actomyosin structure organization, supramolecular fiber organization, and positive regulation of MAPK cascade and signal transduction, reflecting retained contractile and matrix-remodeling identity (**Supplementary Fig. 7e**).

Senescence-like fibroblasts were enriched for regulation of transcription, chromatin remodeling, protein ubiquitination, and protein modification by small protein conjugation (**Supplementary Fig. 7f**), consistent with a senescence-associated stress-responsive regulatory state. Proliferating fibroblasts showed the most divergent GO enrichment profile, dominated by DNA replication, mitotic sister chromatid segregation, mRNA splicing, ribosome biogenesis, and DNA damage response pathways (**Supplementary Fig. 7g**), reflecting active cell cycle progression. Taken together, these subtype-specific transcriptional profiles and pathway enrichments support the biological distinctiveness of cultured fibroblast subtypes identified in the integrated analysis and further reinforce that in vitro culture drives acquisition of proliferative and stress-associated programs alongside partial retention of stromal ECM-remodeling identity.

### In vitro culture drives progressive loss of fibroblast subtype identity and acquisition of inflammatory and senescence transcriptional programs

To characterize the transcriptional state of cultured fibroblasts relative to their lung tissue counterparts, we analyzed cultured fibroblasts across isolation protocols, passages, and sample groups relative to lung tissue fibroblasts from age-matched control, IPF upper lobe, and IPF lower lobe samples. PCA of integrated profiles confirmed that cultured fibroblasts occupied a transcriptional space clearly separated from lung tissue fibroblasts across all conditions and protocols, with progressive divergence observed between initial and final passage (**Fig. 7a**), consistent with accumulating transcriptional drift during prolonged culture. To assess subtype identity retention, we examined the expression of canonical marker genes for each lung tissue fibroblast subtype across isolation protocols and sample types (**Fig. 7b, c; Supplementary Fig. 8**). Alveolar fibroblast markers (alveolar 1 and alveolar 2), including *NPNT*, *INMT*, *RARRES2*, *WNT2*, *CYP4B1*, and *GPC3*, were broadly detectable at initial passage across protocols but progressively attenuated at final passage, with the most pronounced loss observed in WLCS-derived fibroblasts (**Fig. 7b; Supplementary Fig. 8a, b**), suggesting that alveolar subtype identity is not stably maintained in vitro. Adventitial markers including SCARA5 and SFRP2 showed variable retention depending on isolation protocol, while peribronchial and CTHRC1+ subtype markers, notably WIF1, ASPN, CTHRC1 and TGFB1, were undetected across cultured conditions relative to lung sample types (**Fig. 7b**). SMC markers MYH11 and ACTA2 were retained most consistently across passages and protocols.

**Figure 7.**
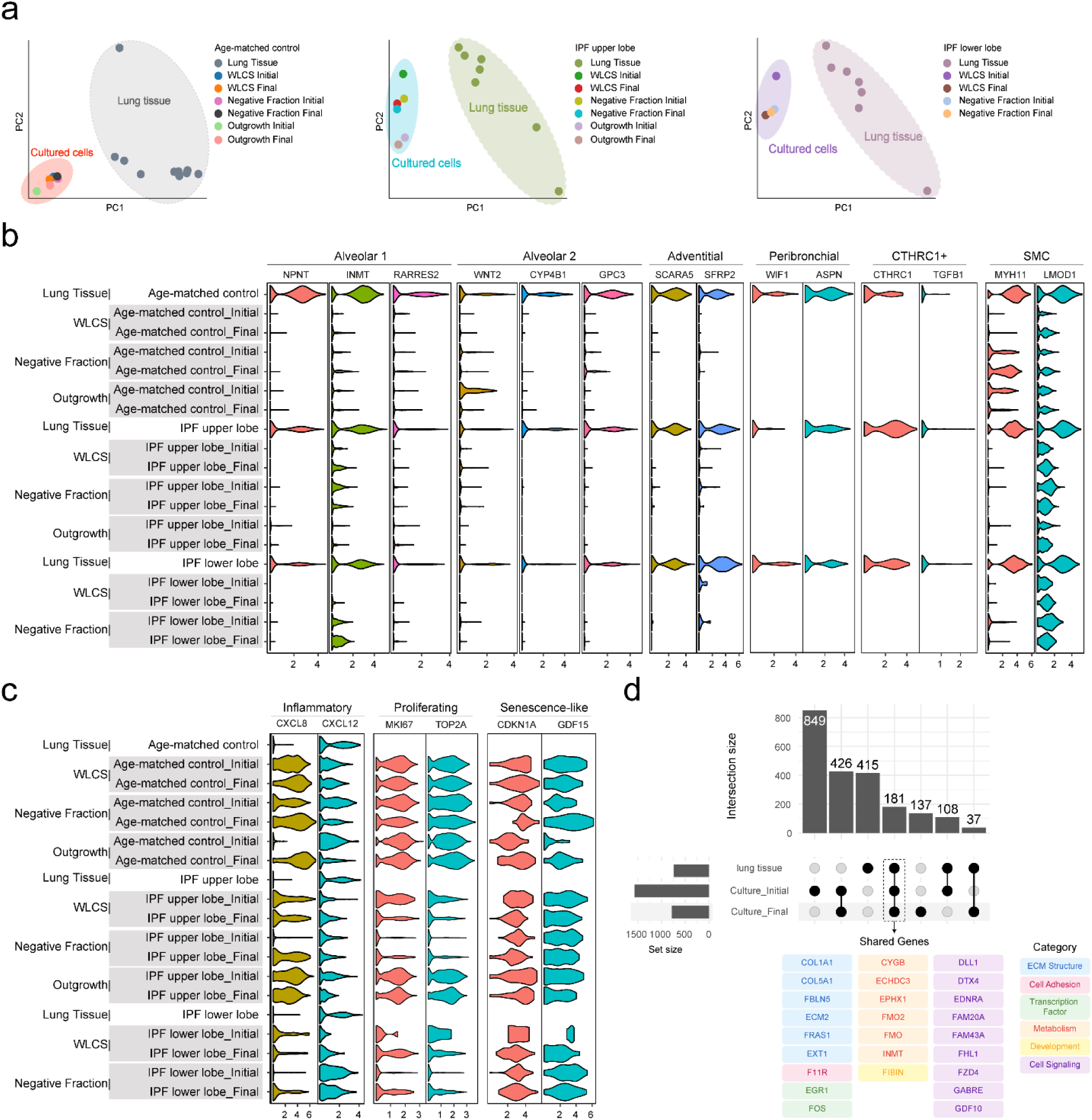
Loss of fibroblast identity and acquisition of inflammatory and senescence signatures during culture. **a.** PCA showing the similarity of gene expression profiles among lung tissue and cultured fibroblast samples. Each color pair represents a distinct protocol. **b.** Violin plots showing decreased expression of lineage-specific markers for alveolar 1, alveolar 2, adventitial, peribronchial, and CTHRC1+ fibroblasts, alongside retained expression of SMC-associated markers and increased expression of inflammatory-state markers. **c.** Violin plots showing elevated expression of proliferative and senescence-like fibroblast markers in cultured cells. **d.** Left: Upset plot illustrating intersections of gene sets from lung tissue and cultured fibroblasts at initial and final passages. Right: Genes shared between lung tissue and cultured fibroblasts across passages, grouped by functional category.

In contrast to the attenuation of tissue subtype markers, cultured fibroblasts progressively acquired expression of inflammatory, proliferating, and senescence-like state markers. Inflammatory markers *CXCL8* and *CXCL12*, proliferation-associated genes *MKI67* and *TOP2A*, and senescence markers *CDKN1A* and *GDF15,* were upregulated at final passage across multiple protocols and conditions (**Fig. 7c**), indicating that extended culture drives acquisition of stress-associated transcriptional programs irrespective of tissue origin. Intersection analysis of DEGs across culture states revealed large culture-specific gene sets, a small shared gene set between lung tissue and cultured fibroblasts (181 genes), with substantial overlap between initial and final cultured states (426 genes), a more limited overlap between lung tissue and cultured fibroblasts, particularly at the last passage (37 shared genes) (**Fig. 7d**), further quantifying the extent of transcriptional remodeling induced by culture. Shared genes spanned ECM structure, transcription factor, metabolism, and cell signaling categories (**Supplementary Table 3**), reflecting retention of broad stromal transcriptional features alongside extensive culture-induced remodeling. Collectively, these findings demonstrate that in vitro culture induces progressive loss of lung fibroblast subtype-specific transcriptional identity, accompanied by acquisition of inflammatory and senescence-associated gene expression programs, with the degree of identity loss varying by subtype and isolation protocol.

### Culture conditions induce broad CTHRC1 upregulation across fibroblast subtypes without formation of a distinct CTHRC1+ cluster

To assess whether the CTHRC1+ fibroblast identity observed in lung tissue is recapitulated in vitro, we examined *CTHRC1* gene expression across cultured fibroblast subtypes at initial and final passages stratified by isolation protocol and disease condition (Fig. 8). In lung samples, *CTHRC1* expression was highest in the CTHRC1+ subtype across all conditions, as expected (**Fig. 8a-c, orange-highlighted points**). However, upon transition to culture, *CTHRC1 gene* expression increased markedly across multiple fibroblast subtypes at initial passage, with the magnitude of induction varying by isolation protocol, and was broadly sustained or further elevated at final passage. This pattern was observed in age-matched control and IPF upper lobe samples across all three isolation protocols and in IPF lower lobe samples across the available WLCS and negative fraction cultures and in IPF lower lobe samples across the available WLCS and negative fraction cultures. Notably, subtypes with low *CTHRC1* expression in lung tissue, including adventitial, alveolar 1, alveolar 2, and peribronchial fibroblasts, acquired substantial CTHRC1 expression upon culture initiation, in some conditions approaching the expression levels observed in the tissue-derived CTHRC1+ subtype. This convergence was observed across all three isolation protocols (WLCS, negative fraction, and outgrowth), although it was strongest in WLCS and outgrowth cultures, suggesting that *CTHRC1* upregulation is a general response to in vitro culture conditions rather than a protocol-specific artifact. Collectively, these findings indicate that culture conditions drive widespread *CTHRC1* transcriptional upregulation across fibroblast subtypes, obscuring the subtype-specific expression pattern characteristic of the tissue CTHRC1+ population and helping explain the absence of a transcriptionally distinct CTHRC1+ cluster in vitro. This further supports the notion that in vitro culture induces transcriptional drift that erodes subtype-specific gene expression signatures.

**Figure 8.**
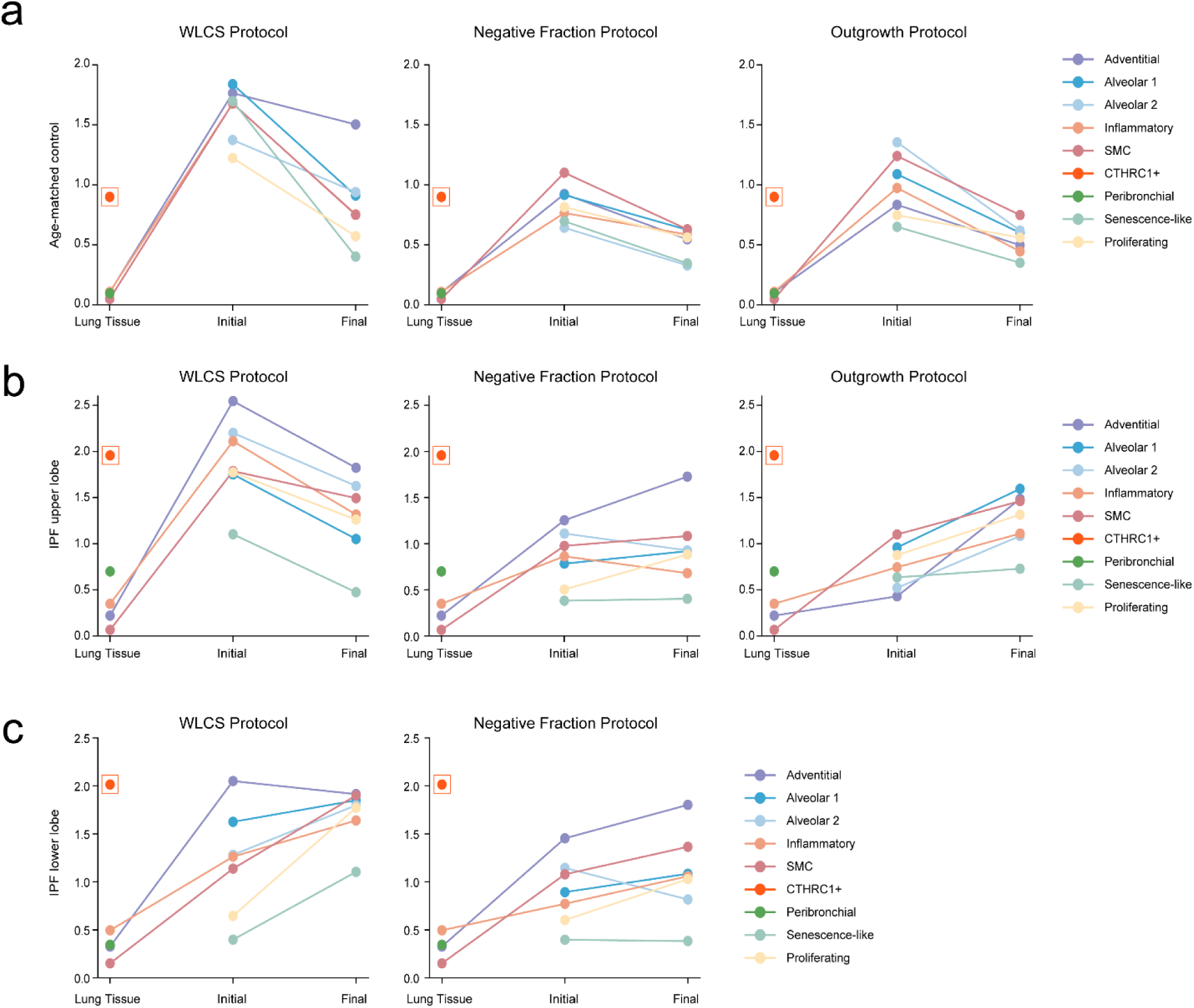
Culture conditions induce CTHRC1 expression across fibroblast subtypes without formation of a distinct CTHRC1+ cluster. Line plots depicting CTHRC1 gene expression levels in lung tissue and cultured fibroblasts at initial and final passages, stratified by fibroblast subtype and isolation protocol. The orange dot enclosed by an orange square denotes high CTHRC1 expression in lung tissue. Colors indicate fibroblast subtypes as shown in the legend. **(a)** Age-matched control samples across WLCS (left), negative fraction (middle), and outgrowth (right) protocols. **(b)** IPF upper lobe samples across WLCS (left), negative fraction (middle), and outgrowth (right) protocols. **(c)** IPF lower lobe samples across WLCS (left) and negative fraction (right) protocols; no outgrowth protocol data are available for this group.

### Transcriptional drift during in vitro culture is constrained by lung fibroblast subtype identity

To investigate the transcriptional relationship between lung tissue fibroblast subtypes and their in vitro cultured counterparts, we integrated scRNA-seq profiles from lung tissue and cultured fibroblasts and applied Waddington Optimal Transport (WOT) to predict subtype-level state transitions. UMAP visualization of the integrated dataset revealed that cultured fibroblasts occupied a transcriptionally distinct region from lung tissue fibroblasts, resolving into seven states including alveolar 1, alveolar 2, proliferating, and senescence-like populations not observed in tissue (**Fig. 9a, b**), indicative of substantial transcriptional drift induced by in vitro culture conditions. WOT-based label transfer onto the tissue fibroblast UMAP confirmed that predicted cultured subtype identities remained spatially coherent with tissue subtype boundaries (**Fig. 9b**), suggesting that drift occurs along subtype-constrained trajectories.

**Figure 9.**
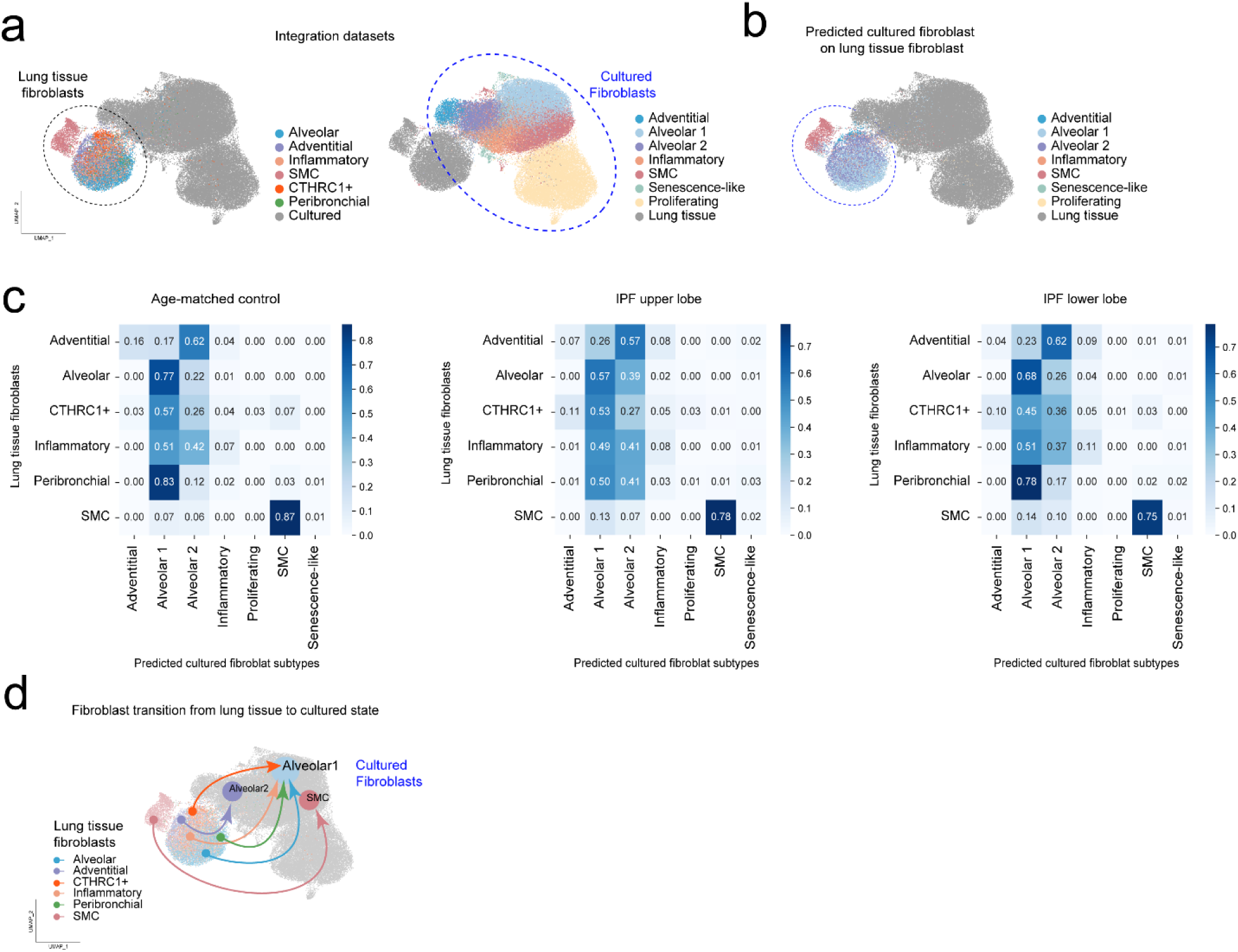
Optimal transport-based prediction of lung tissue fibroblast subtype fate into cultured fibroblast states. **a.** Integrated UMAP embeddings highlighting lung tissue fibroblasts (left, dashed circle) and cultured fibroblasts (right, dashed circle), colored by annotated subtype. Lung tissue fibroblasts include adventitial, alveolar, CTHRC1+, inflammatory, peribronchial, and SMC populations; cultured fibroblasts are additionally resolved into alveolar 1, alveolar 2, proliferating, and senescence-like states. **b.** UMAP of the integrated dataset with lung tissue fibroblasts (dashed circle) colored according to their predicted cultured fibroblast subtype labels, illustrating the transcriptional correspondence between tissue-derived and in vitro states. **c.** Heatmaps displaying the proportion of each lung tissue fibroblast subtype (rows) predicted to correspond to each cultured fibroblast subtype (columns), shown separately for age-matched control, IPF upper lobe, and IPF lower lobe samples. Values represent WOT-derived prediction scores. **d.** UMAP visualization of the integrated dataset overlaid with arrows indicating predicted transitions of lung tissue fibroblast subtypes (color-coded by tissue subtype of origin) toward their predominant in vitro cultured states, primarily alveolar 1, alveolar 2, and SMC.

Quantification of WOT prediction scores across conditions revealed consistent subtype-specific transition patterns despite this drift (**Fig. 9c**). Alveolar fibroblasts showed predominant correspondence to the alveolar 1 cultured state across age-matched control (0.77), IPF upper lobe (0.57) and IPF lower lobe (0.68) samples. Peribronchial fibroblasts were strongly predicted to transition to the alveolar 1 state in age-matched control (0.83) and IPF lower lobe (0.78) samples, while adventitial fibroblasts consistently mapped onto the alveolar 2 cultured state (age-matched control: 0.62; IPF upper lobe: 0.57 and IPF lower lobe: 0.62). SMCs showed the least transcriptional drift, retaining their identity with the highest prediction scores for the SMC cultured state across all conditions (age-matched control: 0.87; IPF upper lobe: 0.78; IPF lower lobe: 0.75). In contrast, CTHRC1+ fibroblasts displayed a more distributed transition pattern across alveolar 1 and alveolar 2 states, suggesting broader susceptibility to transcriptional drift upon culture. Inflammatory fibroblasts likewise showed a distributed transition pattern dominated by alveolar 1 and alveolar 2 cultured states, with only limited correspondence to the cultured inflammatory state.

WOT-derived transport maps further illustrated that lung tissue fibroblast subtypes converged predominantly onto alveolar 1, alveolar 2, and SMC cultured states, with subtype-specific directionality visible in the integrated UMAP (**Fig. 9d**). Collectively, these results indicate that in vitro culture induces pronounced transcriptional drift in lung fibroblast subtypes, with the extent and directionality of drift varying by subtype, most limited in SMCs and most diffuse in CTHRC1+ and inflammatory populations.

## Discussion

This study provides a multiscale transcriptomic characterization of lung fibroblast heterogeneity in human aged healthy and IPF lungs, integrating scRNA-seq and spatial transcriptomics with transcriptional profiling of profiling of cultured primary fibroblasts across multiple isolation protocols and passages. Collectively, our findings resolve the spatial and signaling architecture of disease-relevant subtypes and reveal the transcriptional consequences of in vitro culture, findings with direct implications for the design of translational in vitro IPF models and the identification of candidate therapeutic targets.

Integration of IPF and age-matched control lung scRNA-seq data identified six transcriptionally distinct fibroblast subtypes, alveolar, adventitial, inflammatory, peribronchial, SMC, and CTHRC1+, consistent with prior human IPF atlases.^8,12,19^ Each subtype displayed a characteristic enrichment program: alveolar fibroblasts for ECM organization and cell migration; adventitial for collagen fibril organization and perivascular matrix maintenance; inflammatory for cytokine-mediated signaling; SMC for actomyosin contractile programs; and CTHRC1+ for the broadest pro-fibrotic ECM biosynthesis and MMP signature. CTHRC1+ fibroblasts therefore emerge as the dominant tissue-defined pro-fibrotic subtype in our dataset, with transcriptional, spatial, and intercellular communication features consistent with central roles in collagen-producing and matrix-remodeling; however, functional validation remains required.^8^ In culture, alveolar 2 fibroblasts were in a pro-fibrotic transcriptional state, displaying the clearest activated matrix-producing state with fibrillar collagens and matricellular genes such as THBS2 and SPARC constitutively elevated in IPF fibroblasts and drivers of β-catenin-mediated apoptosis resistance,^20–22^ EFEMP1 (fibulin-3) was a candidate regulator of fibroblast survival and ECM homeostasis across subtypes. GATA6, enriched in pro-fibrotic cultured alveolar 2 fibroblasts, likely reflects TGFβ1-driven ECM programming.^23^ These features suggest that culture can stabilize a pro-fibrotic alveolar-like program even as native tissue-defined subtype identities erode.

A key finding of this study is the spatial enrichment of CTHRC1+ fibroblasts within fibroblastic foci, confirmed by Xenium In Situ spatial transcriptomics. Fibroblastic foci are the defining active lesion of UIP/IPF, representing subepithelial aggregates of myofibroblasts embedded within a pathological ECM and considered sites of ongoing active fibrogenesis from which fibrosis propagates into adjacent alveoli.^24,25^ The spatial enrichment of CTHRC1+ cells within these structures anchors their transcriptional identity, the broadest pro-fibrotic ECM and MMP signature identified in our dataset, to the most disease-relevant histopathological niche in IPF. The co-localization of elevated collagen score and MMP expression score within foci-associated regions supports these structures as sites of concurrent ECM deposition and active-matrix remodeling, not end-stage passive scars.^26^

Notably, our spatial data analyses reveal COL7A1 expression in CTHRC1+-enriched fibroblastic foci, consistent with the recent demonstration that collagen VII protein and transcript are detectable within fibroblastic foci in IPF, with COL7A1 further detected in mesenchymal subsets and upregulated in fibroblasts by TGF-β1 in vitro.^27^ As collagen VII normally anchors stratified epithelia to the interstitial matrix, its ectopic accumulation in fibroblastic foci, alongside expanded basal and aberrant basaloid cell populations, may facilitate the abnormal re-epithelialization that characterizes IPF progression.^27^

CellChat analysis of ligand-receptor interaction networks revealed a disease-stage-dependent reorganization of fibroblast signaling hubs that constitutes a second major finding of this study. In age-matched control lungs, peribronchial and SMCs dominated intercellular communication through Notch– and GPCR/S1P-mediated pathways, consistent with their homeostatic airway and vascular niche roles. Despite their lineage-progenitor status, alveolar fibroblasts were relatively signaling-quiescent in health, consistent with recent murine lineage tracing demonstrating that the alveolar fibroblast lineage, while central to the orchestration of lung inflammation and fibrosis, does so through state transitions rather than constitutive signaling activity.^6^ In established IPF (lower lobe), CTHRC1+ fibroblasts emerged as dominant senders, with collagen, laminin, FN1, and tenascin pathways enriched over age-matched control. The CTHRC1+ sender → inflammatory and alveolar receiver axis is the most disease-relevant unit identified, where sustained ECM and matricellular signaling by CTHRC1+ fibroblasts likely drives transcriptional reprogramming of neighboring fibroblast populations, amplifying the pro-fibrotic niche. The collagen VII-enriched matrix environment of fibroblastic foci^27^ may further modulate the mechanical and biochemical context in which these signaling exchanges occur.

PAGA connectivity and Pseudotime trajectory analyses provide complementary topological evidence supporting the relationship between fibroblast subtypes. Alveolar fibroblasts were consistently positioned at a transcriptional origin, while CTHRC1+ fibroblasts occupied a distal, disease-associated node, a topology consistent with murine lineage tracing evidence that the alveolar fibroblast lineage is the primary cellular source of pathological CTHRC1+ fibroblasts in pulmonary fibrosis.^6–8,28^ The positioning of inflammatory fibroblasts at an intermediate node further suggests that transcriptional reprogramming toward a pro-inflammatory state may precede or accompany the acquisition of the full CTHRC1+ pro-fibrotic program. These analyses are descriptive of transcriptional relatedness and do not establish lineage directionality; experimental validation through lineage tracing or clonal analysis is required before directionality conclusions can be drawn.^29^

A third major finding of this study concerns the in vitro transcriptional behavior of lung fibroblast subtypes. Because lung tissue and cultured fibroblasts form transcriptionally separate clusters incompatible with conventional pseudotime inference, we applied WOT to quantify probabilistic state transitions between tissue and cultured states across fibroblast subtypes, isolation protocols, and disease conditions.^30^ WOT revealed convergent drift toward alveolar 1 and alveolar 2 cultured states across conditions. Quantitatively, SMCs demonstrated the greatest resistance to drift (self-retention scores 0.75-0.87), alveolar fibroblasts showed intermediate retention (0.57-0.77), and CTHRC1+ fibroblasts were the most transcriptionally plastic, with dispersed transition probabilities across multiple cultured states.

Drift patterns were broadly similar between IPF and age-matched control samples, establishing that culture-induced transcriptional remodeling is a general in vitro phenomenon rather than a disease-specific effect. IPF fibroblasts, however, showed greater inter-passage variability, likely reflecting the inherent instability of the pro-fibrotic transcriptional state and selective depletion of CTHRC1+ and senescence-like subpopulations over successive passages. These findings align with and extend recent evidence that lung fibroblasts rapidly adopt transcriptional programs distinct from those observed in vivo upon transfer to standard culture, with identities stabilizing early across passages and cue-based restoration being most effective on freshly isolated cells.^31^The identification of ROCK inhibition in that work as modestly restorative implicates mechanosignaling induced by stiff culture surfaces as a primary driver of identity loss, consistent with the convergent, subtype-independent drift we observed. Extended culture further drove acquisition of inflammatory (*CXCL8, CXCL12*), proliferative (*MKI67, TOP2A*), and senescence-like (*CDKN1A, GDF15*) transcriptional programs across all protocols, reflecting passage-dependent replicative stress independent of disease origin and supporting the conclusion that passage-dependent stress responses are a general feature of cultured fibroblasts rather than a disease-specific property.^31,32^

Culture elevated *CTHRC1* gene expression across all fibroblast subtypes, isolation protocols, and disease conditions, often approaching the levels observed in the tissue-defined CTHRC1+ state. This convergence constitutes a fourth key finding with direct practical implications: it precludes the formation of a transcriptionally distinct CTHRC1+ cluster in vitro and limits the use of standard cultured fibroblasts as models of this disease-relevant population. The subtype-restricted, spatially organized *CTHRC1* expression that defines this subpopulation within fibrotic lung tissue cannot be recapitulated under standard culture conditions, a finding broadly consistent with the rapid loss of native subtype markers documented upon culture transfer^31^. Preserving fibroblast subtype fidelity in vitro will likely require biomechanical conditioning, niche-specific cues, or early post-isolation intervention strategies targeting the mechanosignaling pathways identified as primary drivers of transcriptional drift.

A key advantage of this study design is its longitudinal nature, in which fibroblasts derived from the same donors were profiled at initial (passage 1) and final (passage 6), allowing direct within-donor tracking of cell state transitions and minimizing inter-individual variability as a confounding factor. As with any transcriptomic time-course study, several analytical and biological considerations inform the interpretation of our findings. Although pseudotime and PAGA analyses effectively capture transcriptional topology and intercellular proximity, they do not establish lineage directionality or causal relationships between fibroblast subtypes; resolving true transition directionality would require complementary approaches such as RNA velocity or experimental lineage tracing. Similarly, WOT-derived transition probabilities represent model-based stochastic estimates constrained by the assumption that cell populations evolve along energetically optimal trajectories, a premise that may not hold under the non-physiological pressures of serial passaging, where culture-induced transcriptional drift could introduce alternative or stochastic routes not captured by the optimal transport framework. As such, these transition estimates remain computational inferences pending functional validation. Beyond these analytical constraints, the phenotypic and functional consequences of passage-driven transcriptional drift remain uncharacterized. It is presently unknown whether fibroblasts undergoing culture-induced state transitions retain, acquire, or lose pro-fibrotic effector capacity, exhibit altered responsiveness to antifibrotic or cytotoxic compounds, or engage in aberrant paracrine signaling with neighboring cell populations.

In summary, this study provides an integrated transcriptomic atlas of lung fibroblast heterogeneity in IPF spanning tissue, spatial, and in vitro dimensions. The principal findings are: (i) six transcriptionally distinct fibroblast subtypes with characteristic ECM and signaling enrichment programs, (ii) spatial enrichment of CTHRC1+ fibroblasts within fibroblastic foci coinciding with concurrent collagen deposition, MMP remodeling, and ectopic collagen VII accumulation, (iii) disease-stage-dependent reorganization of signaling hubs in which CTHRC1+ fibroblasts emerge as dominant senders in established IPF, (iv) WOT-quantified subtype-dependent transcriptional drift during culture, with SMCs most resistant and CTHRC1+ fibroblasts most plastic, and (v) culture-wide upregulation of CTHRC1 across all subtypes, precluding faithful in vitro modeling of this population. The alveolar fibroblast lineage occupies a transcriptional origin position consistent with its established role as the progenitor of pathological CTHRC1+ fibroblasts in murine fibrosis models.^6–8^ Collectively, these findings identify spatial niches, ECM components, cell-cell communication axes, and culture-induced transcriptional programs as candidate targets for therapeutic and modeling strategies in IPF.

## Methods

### Human Lung Tissue Collection and Processing

The Ohio State University (OSU) Comprehensive Transplant Center Human Tissue Biorepository (CTC Biorepository) distributed human lung tissue samples from IPF patients and healthy, aged (>65 years) deceased donors. Informed consent under the IRB-approved Total Transplant Care Protocol (2017H0309) was obtained from IPF patients, and upper and lower lobe tissues were collected from the diseased lung after explant during transplant surgery. Research authorization from deceased donors was obtained by our local organ procurement organization, Lifeline of Ohio, and lung tissues were procured for the CTC Biorepository from donation after brain death (DBD) or donation after circulatory death (DCD). Samples were then distributed via an IRB-approved Honest Broker process (20170310) to our laboratory under an IRB-approved secondary research protocol (2021H0180). The David Geffen School of Medicine at the University of California, Los Angeles (UCLA) also provided de-identified healthy donor lung tissue procured from the International Institute for the Advancement of Medicine (IIAM) and de-identified IPF patient explanted diseased lungs procured from the Ronald Reagan UCLA Medical Center during transplant surgery under an IRB-approved protocol (21-000390-CR-00003). Deceased donors had normal lung function and no history of lung disease. A demographic summary of the donor population is shown in **Supplementary Table 1.**

### Primary Human Lung Fibroblasts Isolation and Cell Culture

#### Establishing cultures

Cultures were established by isolating primary lung fibroblasts from lung tissue specimens. Donor characteristics of human primary lung fibroblasts from Non-IPF and IPF subjects are described in **Supplementary Table 2.** Briefly, after removing the visceral pleura, 20 g of lung tissue was washed three times with 1X PBS (Gibco, 10010023) and minced into small pieces. A portion of the lung tissue (5 to 7 g) was transferred to a gentleMACS C-tube (Miltenyi Biotec, 130-093-237) and subjected to mechanical and enzymatic digestion using a gentleMACS Octo Dissociator with Heaters (Miltenyi Biotec, 130-096-427) and a solution that included collagenase type IV (Worthington Biochemical, LS004186), elastase (Worthington Biochemical, LS002292), dispase (StemCell, 7913), and DNase I (Worthington Biochemical, LS002139). After dissociation, 1X cold wash buffer (Dulbecco’s Modified Eagle Medium (DMEM), high glucose, Gibco, 11965118; 1% penicillin, streptomycin, and amphotericin B; 10% fetal bovine serum (FBS), Gibco, 16140071) was used to inactivate enzymes. The digested tissue was filtered through a series of strainers (300-, 100-, 70-, and 30-µm pore sizes), followed by centrifugation and incubation at room temperature with 1X RBC lysis buffer (distilled water, Gibco, 15230162; 10X RBC Lysis Buffer, BioLegend, 420302). The whole lung cell suspension (WLCS) was resuspended in 1X PBS for cell counting and viability assessment by trypan blue. The highest yield of viable cell suspensions was used.

### Fibroblast enrichment and passaging

To obtain fibroblast-enriched single-cell suspensions from the aged and IPF lung tissues, three optimized isolation protocols were used. Protocol 1 consisted of culturing WLCS to select the plastic-adherent population, which was defined as fibroblasts. Non-adherent cells were removed after three days to purify the fibroblast population. Protocol 2 enriched the fibroblast population from WLCS using magnetic bead cell sorting (LS Column, Miltenyi Biotec, 130-042-401) to deplete immune cells (human CD45 microbeads, Miltenyi Biotec, 130-045-801), endothelial cells (CD31 Microbead Kit, human, Miltenyi Biotec, 130-091-935), and epithelial cells (CD326/EPCAM Microbeads, human, Miltenyi Biotec, 130-061-101) to obtain a negative fraction (Neg F) for all antigens. Protocol 3 isolated lung fibroblasts from distal lung tissue cut into 1 cm **×** 1 cm pieces placed in in 6-well plates and submerged in fibroblast medium (DMEM+F12, Gibco, 11330032; 10% FBS, Gibco, 16140071; 1% non-essential amino acids, Gibco, 11140050; 1% GlutaMAX Supplement, Gibco, 35050061) for 3-4 weeks to allow the fibroblasts to “crawl out” of the tissue and form an adherent monolayer. These fibroblast populations were then dissociated with TrypLE Express (ThermoFisher Scientific,12605036). All fibroblasts were cultured in cell culture flasks in a controlled environment at 37°C and 5% CO_2_ in DMEM high glucose (Gibco, 11965118) with 10% FBS (Gibco, 16140071), and 1% penicillin-streptomycin (P/S) (Gibco, 15140122). Culture media were refreshed three times per week to maintain a sterile and optimal environment for cell growth and expansion. Cell adherence and proliferation were monitored, and once cultures reached 80% confluence, cells were passaged using 0.25% trypsin-EDTA (Gibco, 25200056). Fibroblasts were collected at passages 1 (P1 or “initial”) and 6 (P6 or “final”).

### Single-cell RNA-Sequencing

#### scRNA-seq of FFPE lung sections

Human lung tissue was cut into pieces measuring approximately 10 mm **×** 20 mm, fixed in 4% paraformaldehyde for 24 hours, and transferred to 70% ethanol before paraffin embedding to generate formalin-fixed paraffin-embedded (FFPE) tissue blocks. Two to three 10-µm scrolls were obtained from each FFPE block using a microtome (Leica, HistoCore BIOCUT). Total RNA was extracted using the RNeasy DSP FFPE Kit (Qiagen, 73604). RNA quality was assessed using the Distribution Value of fragments >200 nucleotides (DV200 metric), with a threshold of ≥30% applied to determine suitability for successful probe hybridization, library construction, and next-generation sequencing (NGS). Two 50-µm scrolls from each FFPE block were dissociated into a single-cell suspensions. 10x Genomics Demonstrated Protocol CG000632, Rev B and 1 mg/mL of Liberase TH (Millipore Sigma, 5401135001).

#### scRNA-seq of fixed cell suspension

Fibroblasts were fixed using the Chromium Fixed RNA Profiling protocol for fixation of cells and nuclei (10x Genomics, CG000478, Rev D). Briefly, 1-2 **×** 10^6^ fibroblasts with over 90% viability were collected at P1 or P6. The cells were centrifuged at 400 x g for 5 min at 4°C and then the supernatant was discarded. The cell pellet was resuspended in fixation buffer, which included nuclease-free water (ThermoFisher Scientific, AM9937), concentrated fixation and permeabilization buffer (10x Genomics, PN-2000517), and 4% formaldehyde (Fisher Scientific, BP531-25), and cells were incubated for 24 h at 4°C. Following the incubation, cells were centrifuged at 850 x g for 5 min at room temperature, and the supernatant was discarded. The pellet was then resuspended in chilled quenching buffer (10x Genomics, PN2000516), enhancer (10x Genomics, PN-2000482), and glycerol (Millipore Sigma, G5516). Samples were stored at –80°C until scRNA-seq processing. Fixed RNA profiling was performed based on human probes to detect the whole transcriptome and conducted with the Chromium X platform, with a target cell recovery of 128,000 cells. For library construction, DNA was amplified after 10 PCR cycles for indexing (Dual Index Plate TS Set A, Chromium Fixed RNA Profiling Reagent Kits for Multiplexed Samples, 10x Genomics User Guide CG000527, Rev F).

### Sequencing

NGS was carried out by the Advanced Genomics Core at the University of Michigan (UM) and by the Novogene Corporation Inc. Gene expression libraries were sequenced on an Illumina sequencer NovaSeq 6000 with a sequencing depth of 15,000 read pairs per cell, paired-end, dual indexed, with read lengths of 28 cycles Read 1, 10 cycles i7 index, 10 cycles i5 index, and 90 cycles Read 2. Raw sequencing data in FASTQ format were processed and analyzed using Cell Ranger (v7.1.0, 10x Genomics). The pipeline included barcode demultiplexing, alignment to the human reference transcriptome, and quantification of transcripts through Unique Molecular Identifier (UMI) counting. Quality control metrics, including sequencing saturation, read depth per cell, and the proportion of reads mapped to the transcriptome, were assessed. Processed output files were subsequently generated for downstream analyses.

### Xenium In Situ

#### Gene panel design

Xenium In Situ technology relies on a pre-defined gene panel. Each probe consists of two complementary sequences targeting the mRNA of interest and a unique gene-specific barcode. A total of 389 genes were analyzed using Chemistry (version v1). Of these, 289 genes were derived from a Xenium human predesign lung panel by 10x Genomics, while 100 genes were derived from an add-on custom designed CH7CY9 panel by Lung Aging Lab. The custom panel was curated based on human lung single-cell data analysis generated by Lung Aging Lab, focusing on genes relevant for cell type identification and potential involvement in IPF.

#### Cell segmentation

Cell segmentation of Xenium In Situ data was performed using th*e* Xenium In Situ Cell Segmentation Kit (10x Genomics, PN-1000662) was used according to the manufacturer’s instructions. Briefly, tissue sections were stained with the segmentation reagents, enabling precise delineation of individual cell boundaries based on nuclear and cytoplasmic compartment labeling. Segmentation boundaries were defined using the Xenium Onboard Analysis pipeline, which integrates DAPI-based nuclear segmentation with expansion algorithms to approximate cytoplasmic boundaries. Cell-by-feature matrices were generated by assigning transcript counts to individual segmented cells based on their spatial coordinates. Segmentation quality metrics, including cell area distribution, transcript assignment rates, and nuclear overlap scores, were assessed to ensure high-quality single-cell resolution across all tissue sections. Only cells meeting minimum transcript count thresholds were retained for downstream analysis.

#### Sample preparation

FFPE lung tissue sections (5 mm × 6 mm, 5 µm thickness) were mounted onto Xenium slides and processed according to the 10x Genomics User Guide CG000749, Rev A. Sections underwent deparaffinization and RNA retrieval, followed by hybridization with circularizable DNA probes. Probe ligation generated circular DNA templates that were subsequently amplified by rolling circle amplification, producing high-intensity fluorescent signals with high signal-to-noise ratio. Slides were loaded onto the Xenium Analyzer (Serial # XETG00239) and subjected to iterative cycles of fluorescent probe hybridization, imaging, and signal removal, generating a unique optical signature for each target gene, thereby enabling single-molecule transcript identification at subcellular resolution. Transcript detection, cell segmentation, and spatial mapping were performed using Xenium Onboard Analysis software (v2.0.0.10) and instrument software (v2.0.1.0), comprising of comprehensive onboard analysis for real-time transcript decoding and cell segmentation using the Xenium In Situ Cell Segmentation Kit (10x Genomics, PN-1000662), and off-instrument analysis leveraging the exploration-ready output for downstream interpretation. Final transcript-to-cell assignment, spatial transcript localization, and segmentation quality control were performed in Xenium Explorer v4.0 (10x Genomics), enabling subcellular visualization and interactive exploration of the spatial transcriptomic data across all tissue sections.

### Bioinformatics Analysis

#### Preprocessing

scRNA-seq data were processed in Python (v3.10.8) using Scanpy (v1.11.3).^33^ A random seed of 42 was used to ensure reproducibility. Quality control was applied by retaining cells with a mitochondrial gene fraction below 10% and between 300 and 8,000 detected genes per sample. Potential doublets were removed using scDblFinder (v1.16.0)^34^ in R (v4.3.3) under default parameters, and only singlet cells were retained for subsequent analysis. Following quality control, count matrices from all samples were merged into two separate objects: one for lung tissue cells and another for cultured cells. Library size normalization was performed by scaling each cell to a total of 10,000 counts, followed by log2 transformation of the normalized expression values. The top 3,000 highly variable genes were identified per sample and consolidated for downstream analysis. To mitigate technical variation, gene expression values were regressed against total UMI counts and the percentage of mitochondrial gene counts. The residuals were subsequently scaled, with the maximum scaled value for each gene capped at 10. Principal component analysis (PCA) was performed on the scaled data using the “arpack” solver, and batch effects were corrected by applying Harmony (v0.0.10)^35^ to the resulting principal components (PCs). To accommodate the distinct scale and complexity of each dataset, k-nearest neighbor graphs were constructed separately: for lung tissue cells, using the first 10 Harmony-corrected PCs with 20 neighbors, and for cultured cells, using the first 20 Harmony-corrected PCs with 30 neighbors.

#### Lineage annotation

Lineage-level visualization for lung tissue cells was generated using UMAP.^36^ Unsupervised clustering was then performed on each graph across a series of resolution parameters using the Leiden algorithm (leidenalg v0.10.1)^37^ to define initial cell populations. The resulting clusters were annotated into major lineages based on established canonical marker genes: epithelial cells (*EPCAM, CDH1*), mesenchymal cells (*COL1A1, PDGFRA*) and negative for *PTPRC, EPCAM, PECAM1*, immune cells (*PTPRC, CPA3, CD3D, CD68, JCHAIN, CD79A*), and endothelial cells (*PECAM1, CDH5*) and negative for *PTPRC*. Annotation was supported by marker gene expression heatmaps.

#### Subtype annotation

*T*o enable a more granular annotation of mesenchymal cell subtypes, a second round of analysis was conducted for both lung tissue cells and cultured cells. The mesenchymal populations identified in the first round were subsetted and re-analyzed independently. The full preprocessing and batch correction pipeline, including steps from highly variable gene selection through Harmony integration, was applied to each subset. Subsequently, k-nearest neighbor graphs were constructed with optimized parameters: lung tissue mesenchymal cells were analyzed using 5 neighbors and the first 20 Harmony-corrected PCs, while cultured mesenchymal cells were analyzed using 10 neighbors and the first 20 Harmony-corrected PCs. Subtype-level visualizations were generated for lung tissue mesenchymal and cultured mesenchymal cells separately using UMAP. Unsupervised clustering was then performed on each graph across a series of resolution parameters using the Leiden algorithm to define fibroblast subpopulations. These subclusters were annotated into specific subtypes based on marker genes listed below.^6–8,19,38^ Alveolar, *NPNT, CES1, WNT2, TCF21, INMT, FIBIN, AOC3, GPM6B, SCN7A, FMO2, CYP4B1, LIMCH1, FGFR4, ITGA8, RARRES2, GPC3*; adventitial, *PI16, MFAP5, SCARA5, PTGIS, IGFBP6, SFRP2*; inflammatory, *CXCL2, CXCL8, CCL2, CXCL12, HAS2, PTGDS*; SMC, *MYH11, MYL9, LMOD1, ACTA2;* CTHRC1+, *CTHRC1, POSTN, TGFB1, COL10A1*; peribronchial, *FGF18, WIF1, ASPN* with annotation supported by marker gene expression heatmaps (Fig. 1f).

### Differential Gene Expression and pathway enrichment analysis

Differential gene expression (DGE) was performed within each subtype using the Wilcoxon rank-sum test to compare groups or conditions. Significant genes were defined as those with |log_2_ (fold change) | > 1 and adjusted p-value < 0.05 and were visualized using volcano plots. For each fibroblast subtype, genes used for pathway enrichment were selected using log_2_(fold change) > 0.5 and adjusted p-value < 0.05 from two comparisons: IPF upper lobe vs. age-matched control and IPF lower lobe vs. age-matched control. Enrichment analysis was conducted using Enrichr via the GSEApy Python package (v1.1.9)^39^ against the Gene Ontology Biological Process database.^40^ Dot plots display seven selected pathways, with gene ratio and false discovery rate (FDR) shown for each pathway.

### Trajectory and transcription factor activity analysis

Partition-based graph abstraction (PAGA)^29^ was first constructed to quantify the connectivity among all fibroblast subtypes. Pseudotime trajectories were inferred separately for each group or condition using a Python implementation of Monocle3^41^ on the alveolar, inflammatory, and CTHRC1+ fibroblast populations. Transcription factor (TF) activity was estimated for each cell using decoupler (v 2.1.1)^42^ with the Univariate Linear Model (ULM) method and a curated TF-target interaction resource, CollecTRI.^43^ For each group, TFs were ranked according to the distance correlation between inferred TF activity and pseudotime using the decoupler.tl.rankby_order function. The top seven TFs with the strongest correlations were selected. Temporal changes in TF activity along pseudotime were visualized using line plots, and the spatial distribution of TF activity across cells was additionally displayed on UMAP embeddings.

### Transcriptional drift analysis

Lung tissue and cultured fibroblasts were integrated with scvi-tools^44^ to learn a shared latent embedding across conditions. Transcriptional drift from lung tissue to the cultured state was then evaluated using WOT^30^ by estimating transition probabilities from lung tissue fibroblasts to cultured cells and using these probabilities to predict cultured annotations for lung tissue cells. For each lung tissue annotation, proportion heatmaps summarize the predicted drift patterns across cultured annotations.

### Cell-cell communication analysis

Cell-cell communication (CCC) among fibroblast subtypes was analyzed using CellChat.^45^ Lung tissue fibroblasts were stratified into three groups, age-matched control, IPF upper lobe, and IPF lower lobe, and CellChat was run separately for each group to compare communication patterns across groups. Differential analysis was then performed to identify group-specific changes in CCC between IPF upper lobe and age-matched control, and between IPF lower lobe and age-matched control.

### Statistical analysis

Differential gene expression analysis was performed using the Wilcoxon rank-sum test with Benjamini-Hochberg correction for multiple testing. Pathway enrichment analysis was performed using Fisher’s exact test with Benjamini-Hochberg correction. Differences in collagen family scores between groups and conditions within each fibroblast subtype were assessed using the Wilcoxon rank-sum test. For pseudotime analysis, the top seven TFs in each group were ranked according to their association with pseudotime using distance correlation t-test from dcor^46^, which tests for statistical dependence between TF activity and pseudotime.

## Acknowledgements

We thank the Human Tissue Biorepository at the OSU Wexner Medical Center Comprehensive Transplant Center, The Ohio State University for providing human lung tissue samples used in this study. We also wish to acknowledge the patients and their families for their generous donation of tissue for research purposes.

## Author contributions

This study was designed by M.R., D.T., N.V. Collected data by N.V., V.P., C.S., which was analyzed by N.V., H.C., A.G., J.R-L. and interpreted by N.V., H.C., M.R., J.W., D.T., A.L.M. Additional expertise was contributed by K.S.

The manuscript was drafted by N.V., and revised by B.C., M.R., D.T., A.L.M, B.G, Q.M

## Additional information

### Supplementary Information

**Supplementary Figure 1.**
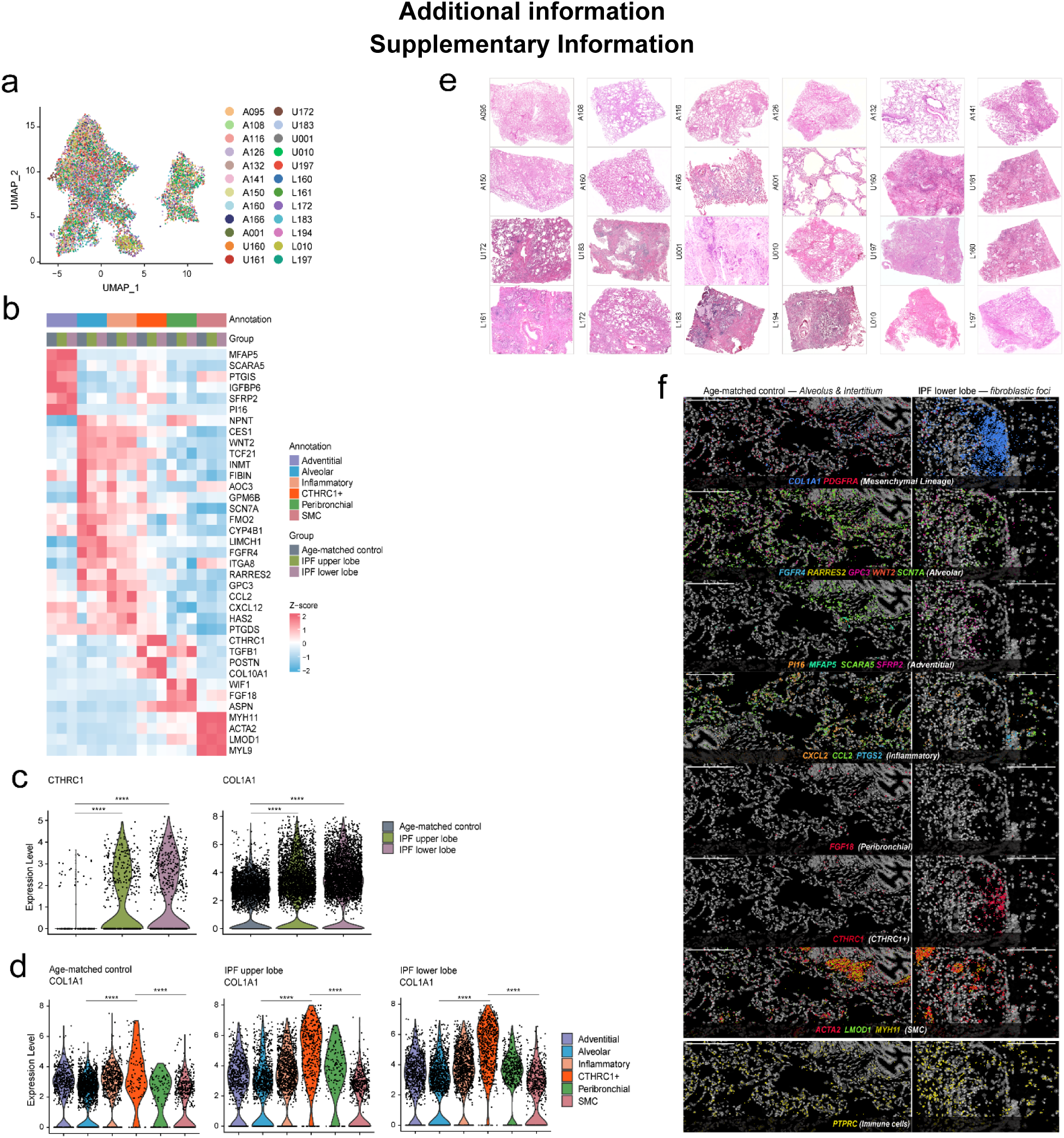
Multimodal visualization of histologic regions and spatial marker expression in IPF lungs. **a.** UMAP visualization shows of mesenchymal cells colored by sample of origin from aged donor lungs and IPF explant lungs. **b.** Heat map of Z-scored expression for selected marker genes across annotated mesenchymal populations. Top bars indicate cell-type annotations and sample origins (age-matched control, IPF upper lobe, IPF lower lobe). Red box highlights a gene cluster with coordinated variation. **c.** Violin plots showing single-cell expression of CTHRC1 and COL1A1 across age-matched control, IPF upper lobe, and IPF lower lobe. Points represent individual cells. Significance is indicated as *p < 0.05, ***p < 0.001, ****p < 0.0001, and ns, not significant. **d.** Violin plots of COL1A1 expression across annotated cell populations in age-matched control (top), IPF upper lobe (Middle), and IPF lower lobe (bottom). Colors denote cell-type annotations; points show single-cell values. Significance is indicated as *p < 0.05, ***p < 0.001, ****p < 0.0001, and ns, not significant. **e.** Histology images of corresponding samples stained with hematoxylin and eosin (H&E) illustrating tissue architecture of all samples used for scRNA-seq. **f.** Xenium In Situ images showing spatial distribution of mesenchymal gene markers in age-matched control tissue and IPF fibroblastic foci. Scale bar, 200 μm.

**Supplementary Figure 2.**
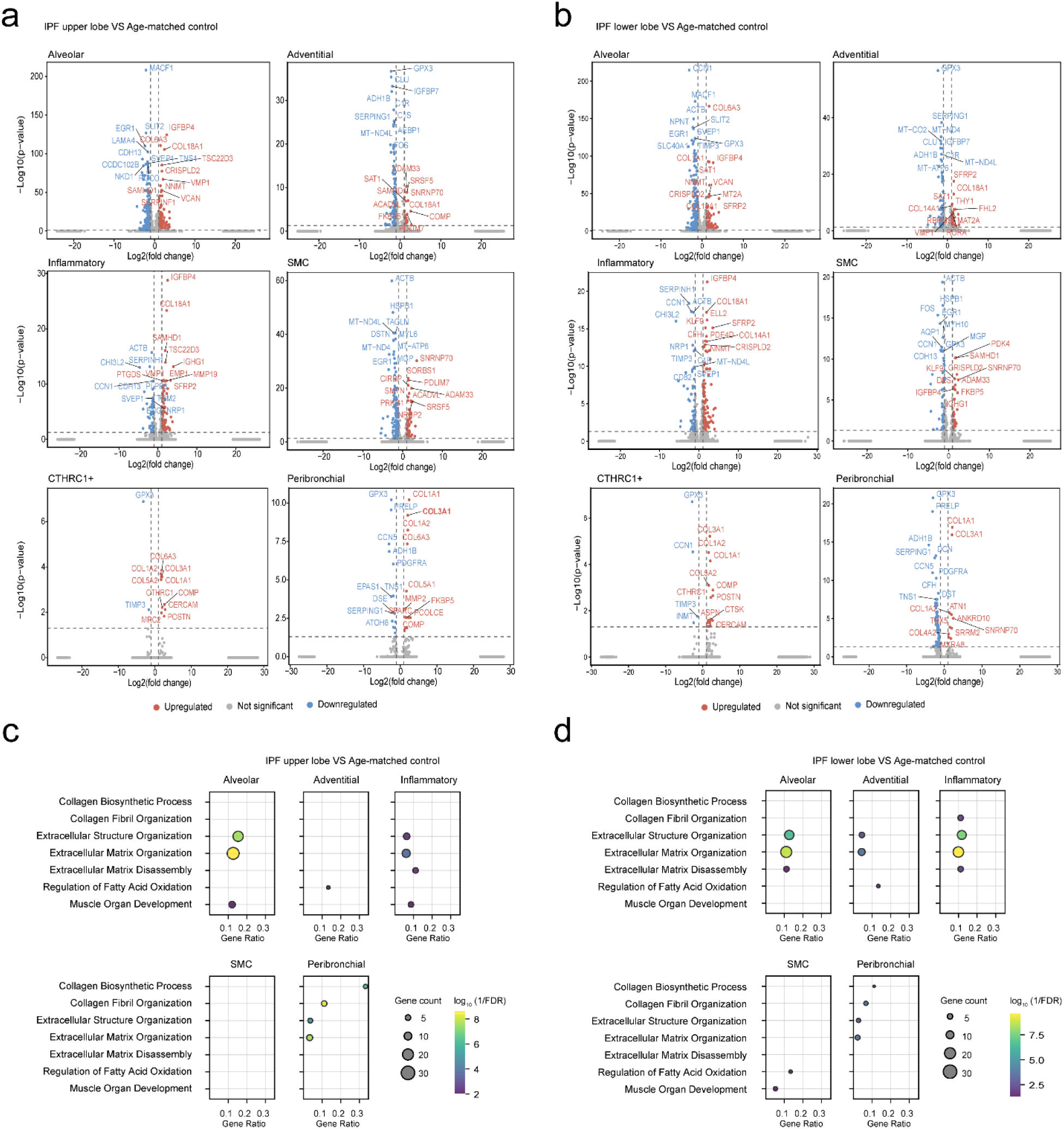
Differential Gene Expression and Fibrosis-Related Pathway Enrichment Across Lung Tissue-Derived Mesenchymal Subtypes. **a-b**. Volcano plots show differentially expressed genes between mesenchymal subtypes from age-matched control tissue versus IPF upper lobe **(a)** and IPF lower lobe **(b)**. Significantly upregulated genes are highlighted in red and significantly downregulated genes in blue. The x-axes represent log₂ fold change and y-axes represent –log₁₀(p-value), with higher values indicating greater statistical significance. **c-d.** Functional pathway analyses of mesenchymal subtypes comparing age-matched control samples versus IPF upper lobe **(c)** or age-matched control samples versus IPF lower lobe samples **(d)**.

**Supplementary Figure 3.**
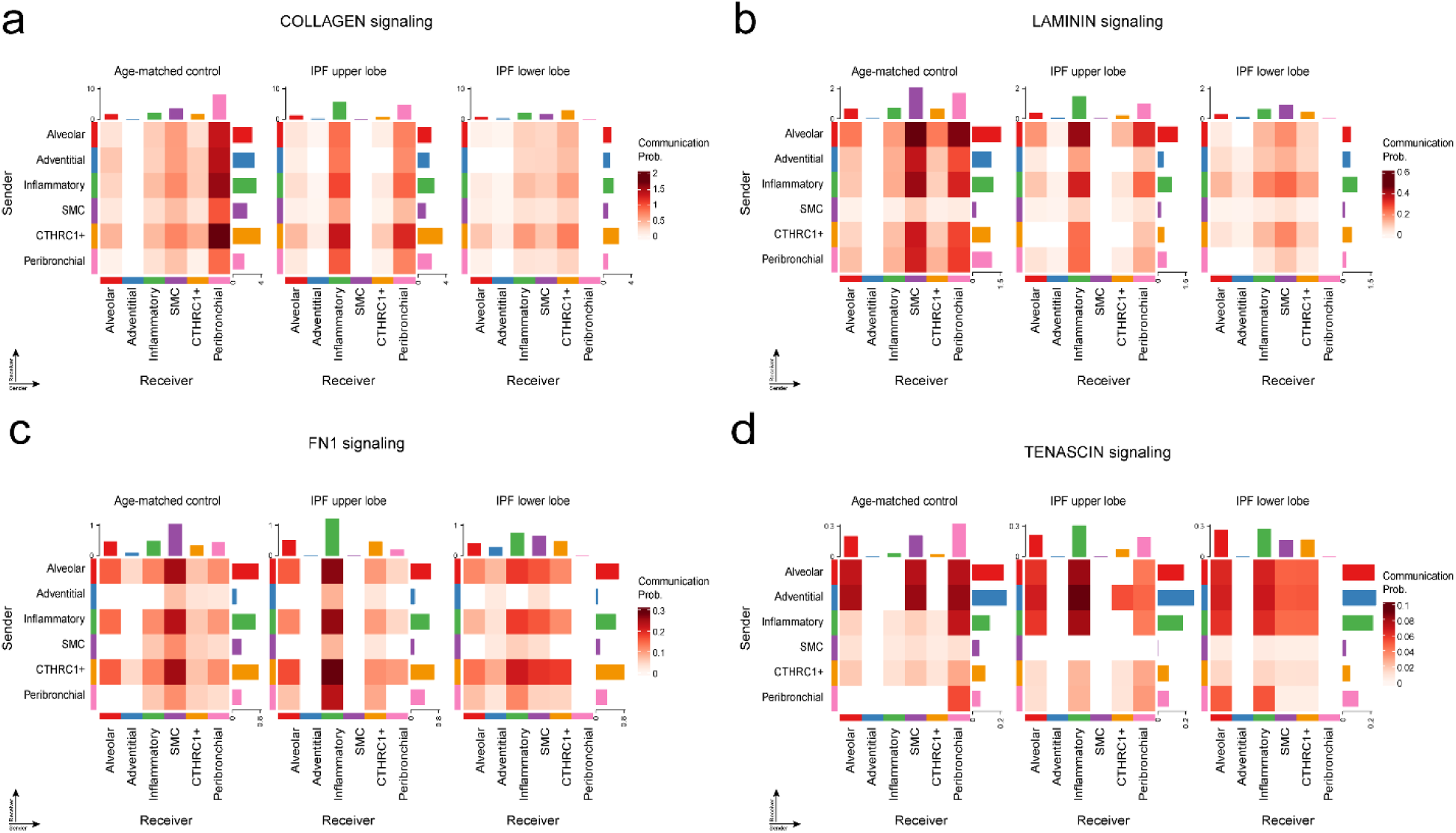
Cell–cell communication probabilities for ECM-related signaling pathways in IPF lung tissue. Heatmaps show predicted ligand–receptor interaction probabilities for four extracellular matrix signaling pathways— collagen **(a)**, laminin **(b)**, fibronectin (FN1) **(c)**, and tenascin **(d)**—across lung tissue–derived mesenchymal subtypes in age-matched control, IPF upper-lobe, and IPF lower-lobe samples. For each pathway, sender (y-axis) and receiver (x-axis) cell groups include alveolar, adventitial, inflammatory, CTHRC1⁺, and peribronchial fibroblasts and SMCs. Color intensity represents the predicted communication probability, as indicated by the accompanying scale bars. Each panel displays pathway-specific interaction strengths within and between cell populations, highlighting context-dependent signaling patterns across disease states and anatomical regions.

**Supplementary Figure 4.**
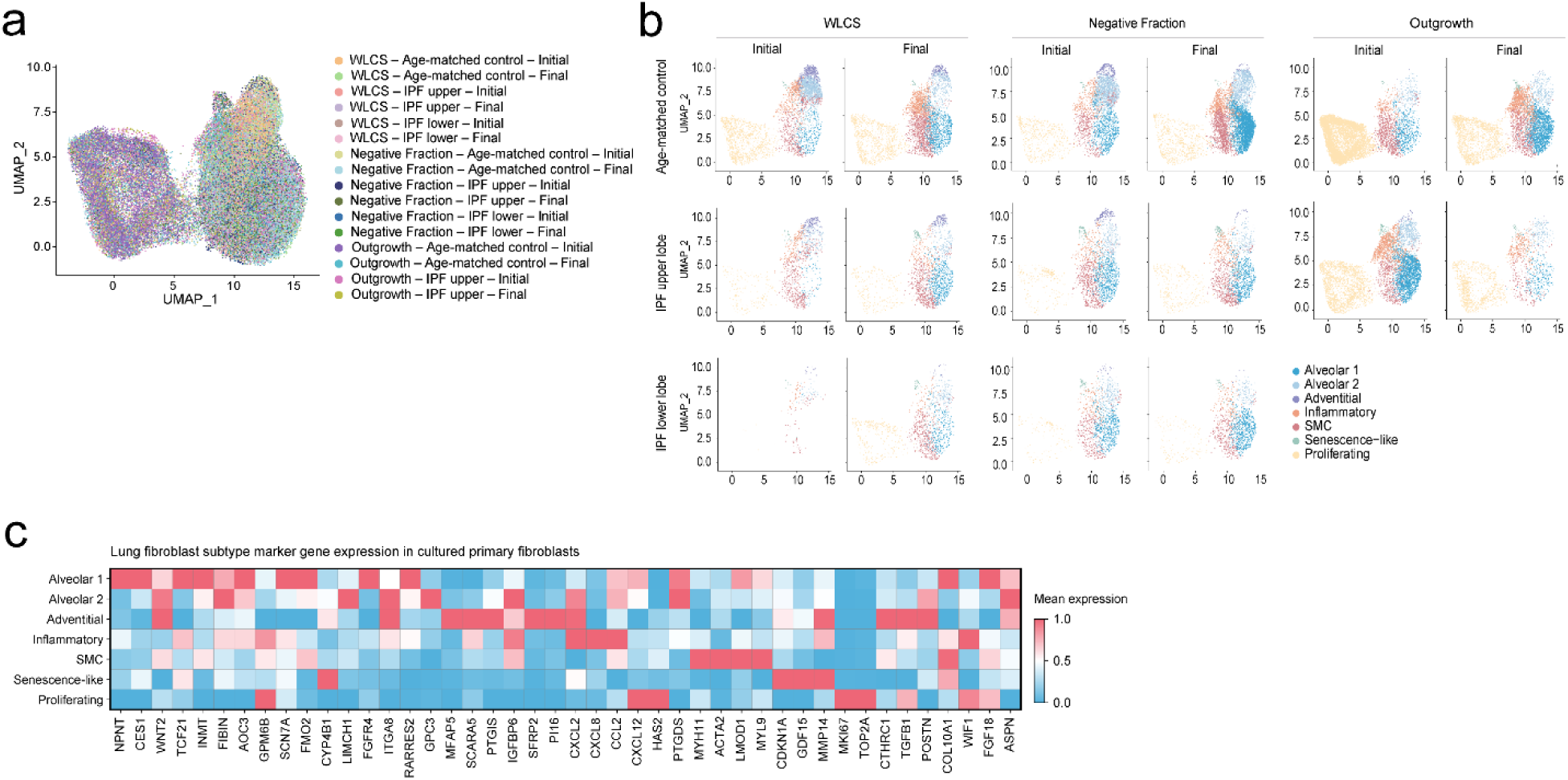
UMAP visualization and gene expression profiles of fibroblast subtypes across isolation protocols. **a.** UMAP plot showing clustering of fibroblast subtypes isolated from all age-matched control and IPF lung samples, colored by sample of origin: alveolar 1, alveolar 2, adventitial, inflammatory, SMC, senescence-like, and proliferating fibroblasts. **b.** UMAP plots stratified by isolation protocol (WLCS, negative fraction, and outgrowth), illustrating the distribution of fibroblast subtypes across isolation protocols, sample groups, and passages. **c.** Heat map displaying mean marker gene expression across fibroblast subtypes, highlighting differences in gene signatures among alveolar 1, alveolar 2, adventitial, inflammatory, SMC, senescence-like, and proliferating populations.

**Supplementary Figure 5.**
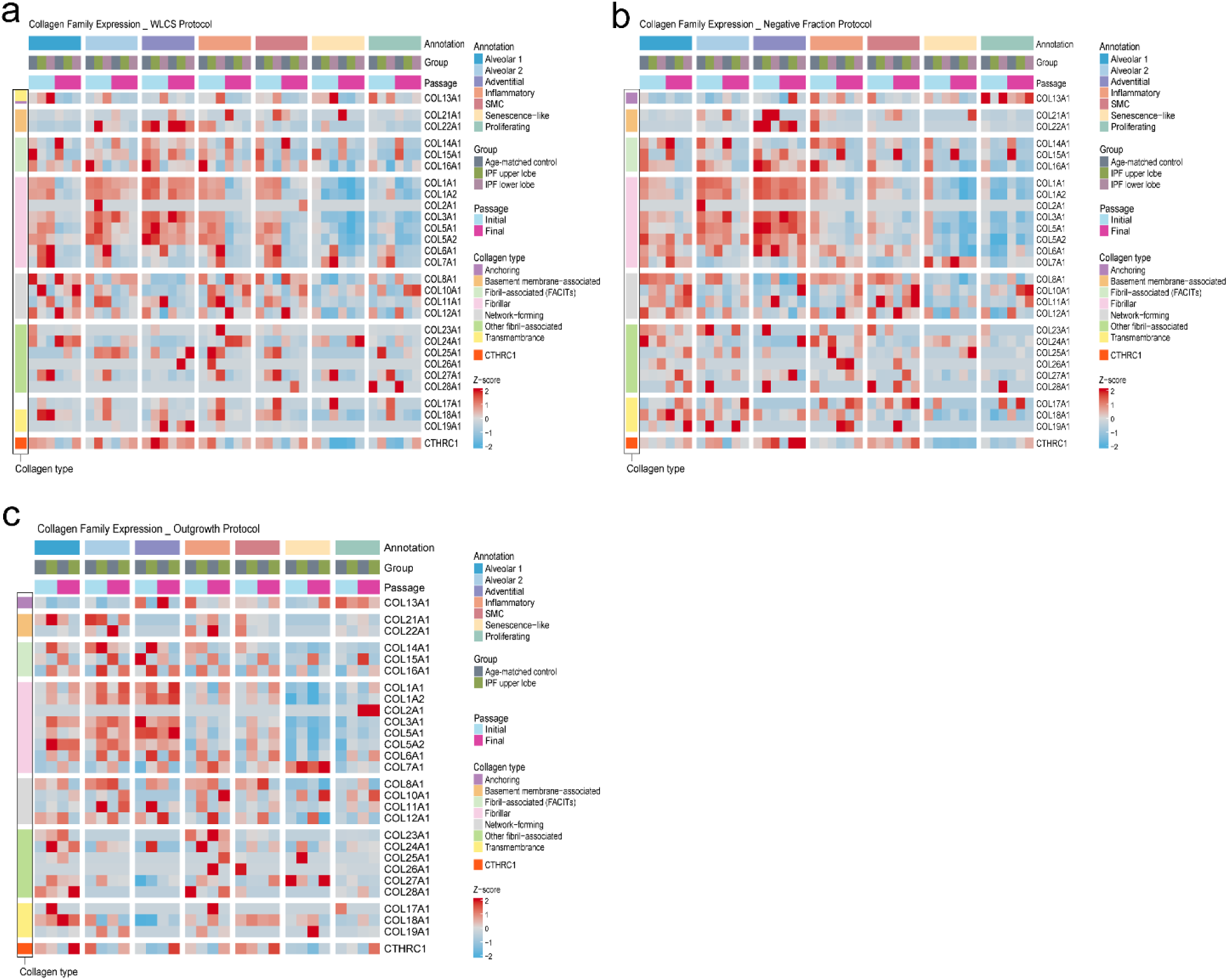
Passage– and protocol-dependent collagen family expression in fibroblast subtypes. **a-c**. Heatmap of collagen family gene expression in cells isolated using three isolation protocols: WLCS (**a)**, negative fraction **(b)**, or outgrowth **(c)**. Top bars indicate subtype annotation, condition, and passage. Collagen family categories are annotated on the left, and Z-score scaling indicates relative expression levels. Outgrowth cultures were available for age-matched control and IPF upper lobe samples only.

**Supplementary Figure 6.**
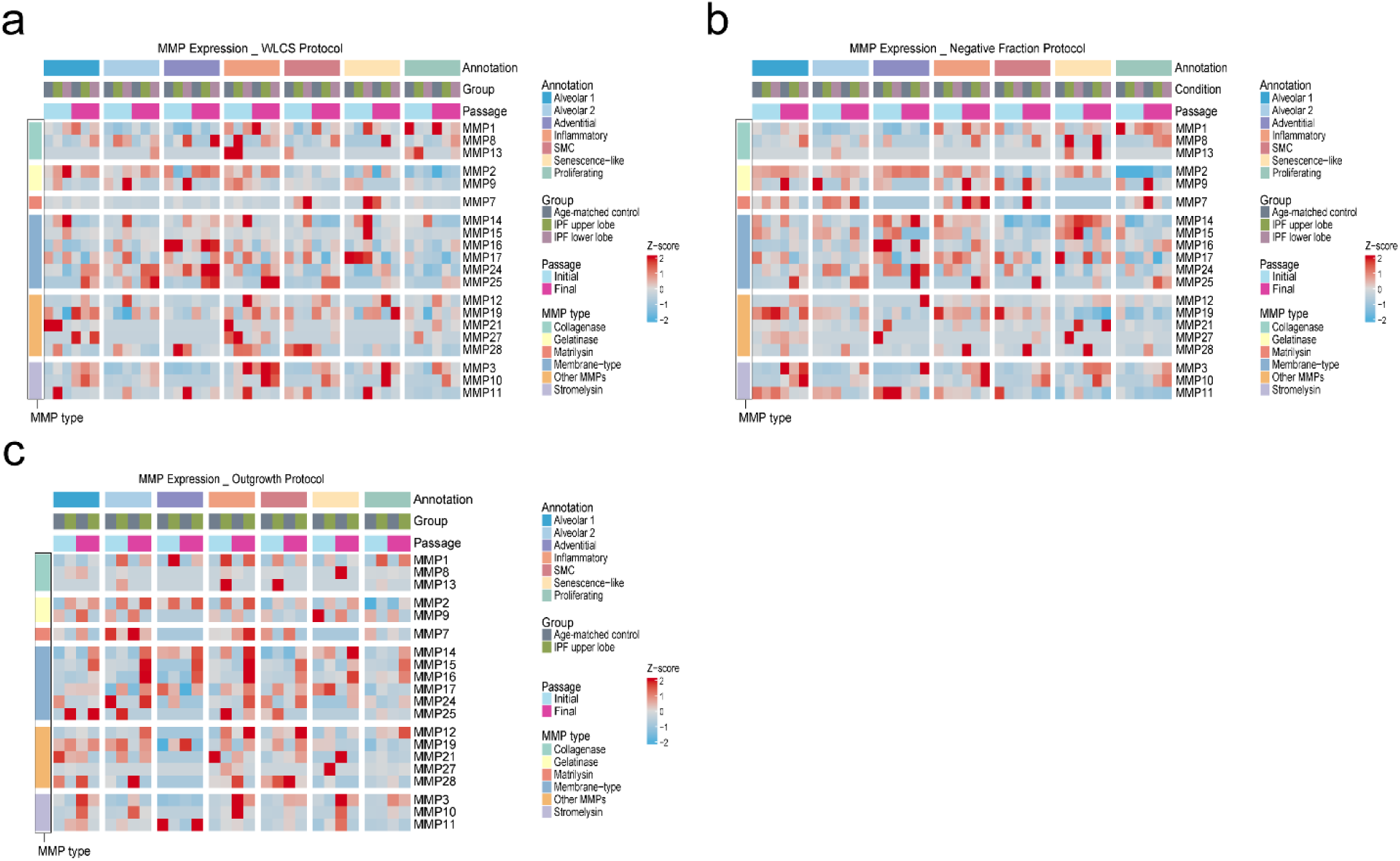
MMP expression across expansion protocols. **a-c**. Z-score-scaled expression of MMP1-MMP28 across annotated lung-derived fibroblast subtypes (alveolar 1, alveolar 2, adventitial, inflammatory, SMC, senescence-like, proliferating) using WLCS **(a),** negative fraction **(b)**, or outgrowth **(c)** isolation protocols from age-matched control, IPF upper lobe, and IPF lower lobe tissue. Top bars indicate subtype annotation, sample group, and passage, with columns including initial and final passage states. MMPs are grouped by functional class; color intensity reflects relative expression.

**Supplementary Figure 7.**
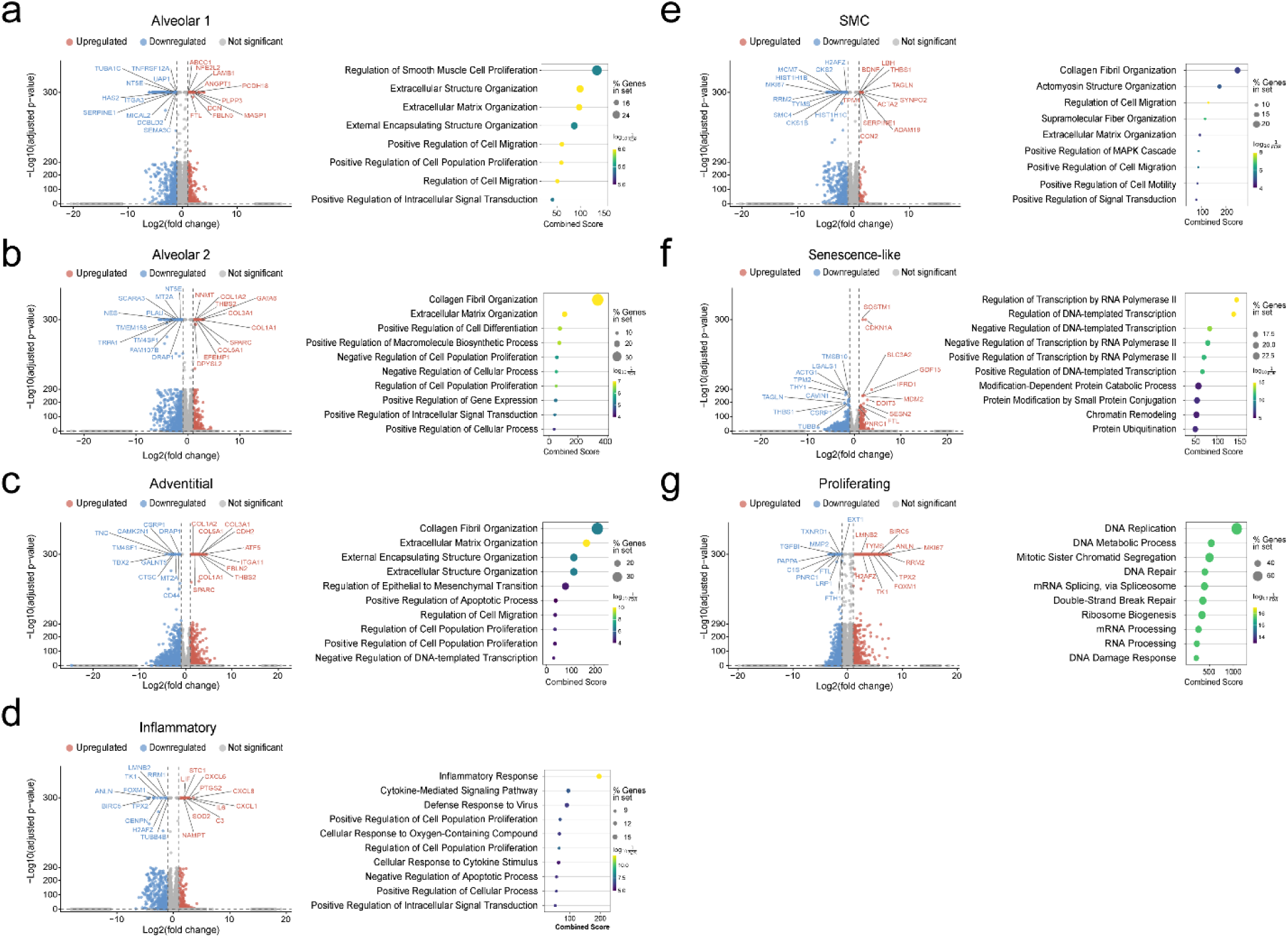
Differential gene expression and pathway enrichment across cultured fibroblast subtypes. Volcano plots of fibroblast subtypes, including alveolar 1 **(a)**, alveolar 2 **(b),** adventitial **(c)**, **i**nflammatory **(d)**, SMC **(e)**, senescence-like **(f),** and proliferating **(g)**.Volcano plots display log₂(fold change) on the x-axis and –log₁₀(FDR) on the y-axis, with upregulated genes in red and downregulated genes in blue. The adjacent dot plot shows significantly enriched GO biological processes ranked by combined score. One-versus-rest differential expression by fibroblast subtype across all isolation protocols, passages, and sample types.

**Supplementary Figure 8.**
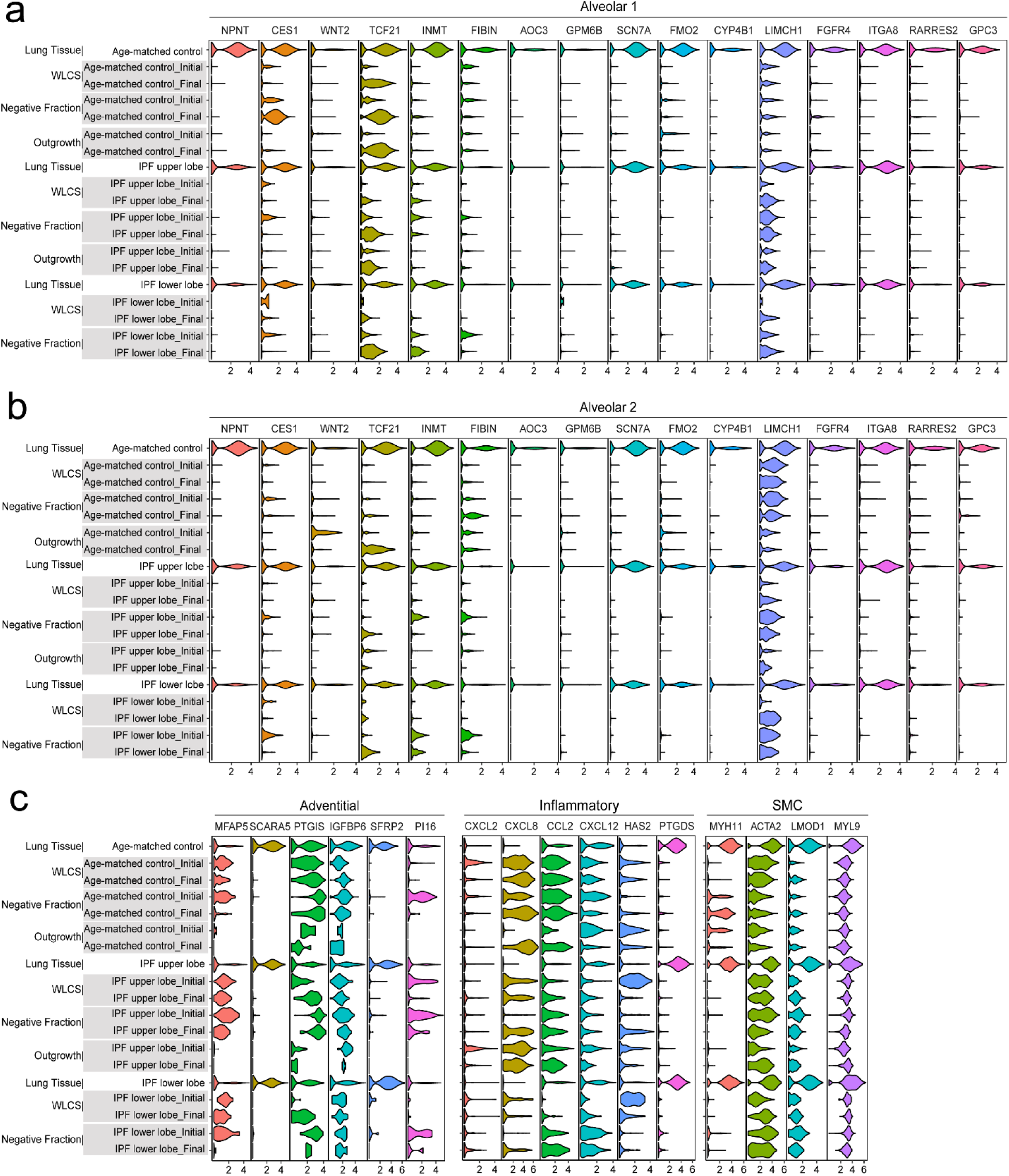
Fibroblast subtype-specific marker gene expression in cultured fibroblasts from age-matched control and IPF lungs. Violin plots showing expression of specific sets of fibroblast subtype marker genes in cultured fibroblasts from age-matched control and IPF samples compared with fibroblasts directly isolated from fixed lung tissue. **a.** Alveolar fibroblasts-1: NPNT, CES1, WNT2, TCF21, INMT, FIBIN, AOC3, GPM6B, FMO2, CYP4B1, LIMCH1, FGFR4, ITGA8, RARRES2, GPC3. **b.** Alveolar-2 fibroblasts: NPNT, CES1, WNT2, TCF21, INMT, FIBIN, AOC3, GPM6B, SCN7A, FMO2, CYP4B1, LIMCH1, FGFR4, ITGA8, RARRES2, GPC3. **c.** Adventitial fibroblasts: MFAP5, SCARA5, PTGIS, IGFBP5, SFRP2, PI16; inflammatory fibroblasts: CXCL2, CXCL8, CCL2, CXCL12, HAS2, PTGDS; SMC: MYH11, ACTA2, LMOD1, MYL9.

**Supplementary Table 1.** Demographic characteristics of lung tissue samples from Non-IPF donors and IPF patients.

| **Sample ID** | **Donor type/Patient** | **Group** | **Lobe** | **Anatomical Region** | **Age** | **Sex** | **Race** | **Smoking history** |
| --- | --- | --- | --- | --- | --- | --- | --- | --- |
| A095 | DCD | Aged-matched control | lower | Parenchyma | 58 | Male | White | Yes |
| A108 | DCD | Aged-matched control | lower | Parenchyma | 61 | Female | White | Yes |
| A116 | DCD | Aged-matched control | lower | Parenchyma | 62 | Male | Afro-American | No |
| A126 | DCD | Aged-matched control | lower | Parenchyma | 61 | Female | White | No |
| A132 | BD | Aged-matched control | lower | Parenchyma | 72 | Female | White | No |
| A141 | BD | Aged-matched control | lower | Parenchyma | 65 | Male | White | Yes |
| A150 | BD | Aged-matched control | lower | Parenchyma | 71 | Female | White | No |
| A160 | DCD | Aged-matched control | lower | Parenchyma | 60 | Male | White | Yes |
| A166 | DCD | Aged-matched control | lower | Parenchyma | 66 | Male | White | No |
| A001 |  | Aged-matched control | lower | Parenchyma | 66 | Female | White | No |
| U160 | IPF Explant | IPF upper | Upper | Parenchyma | 67 | Female | White | Yes |
| U161 | IPF Explant | IPF upper | Upper | Parenchyma | 55 | Male | White | Yes |
| U172 | IPF Explant | IPF upper | Upper | Parenchyma | 59 | Male | White | Yes |
| U183 | IPF Explant | IPF upper | Upper | Parenchyma | 71 | Male | White | Yes |
| U197 | IPF Explant | IPF upper | Upper | Parenchyma | 69 | Male | White | No |
| U001 | IPF Explant | IPF upper | Upper | Parenchyma | 70 | Male | White | Yes |
| U010 | IPF Explant | IPF upper | Upper | Parenchyma | 69 | Female | White | No |
| L160 | IPF Explant | IPF lower | Lower | Parenchyma | 67 | Female | White | Yes |
| L161 | IPF Explant | IPF lower | Lower | Parenchyma | 55 | Male | White | Yes |
| L172 | IPF Explant | IPF lower | Lower | Parenchyma | 59 | Male | White | Yes |
| L183 | IPF Explant | IPF lower | Lower | Parenchyma | 71 | Male | White | Yes |
| L197 | IPF Explant | IPF lower | Lower | Parenchyma | 69 | Male | White | No |
| L010 | IPF Explant | IPF lower | Lower | Parenchyma | 69 | Female | White | No |
| L194 | IPF Explant | IPF lower | Lower | Parenchyma | 70 | Male | White | No |

**Supplementary Table 2.**
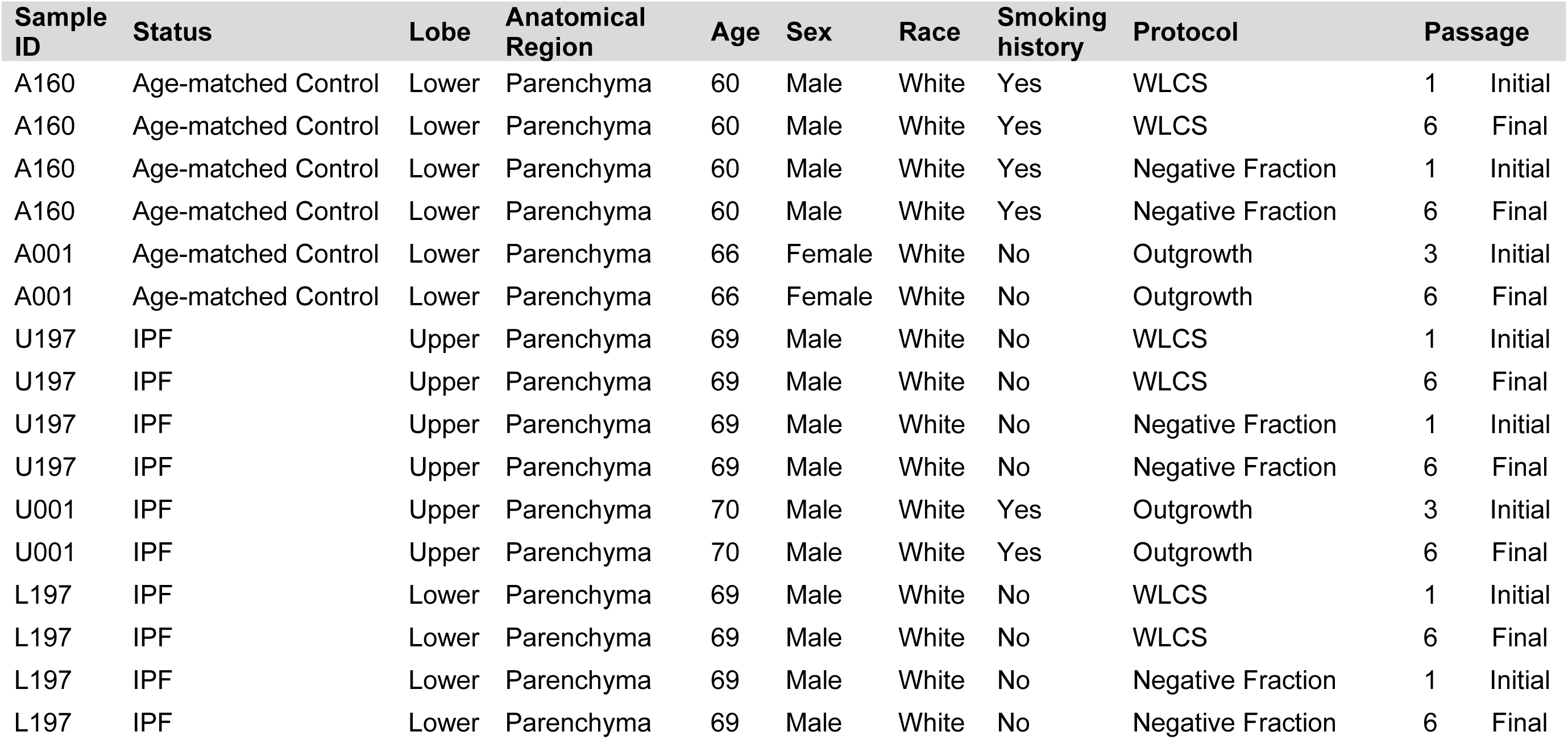
Donor characteristics of human primary lung fibroblasts from Non-IPF and IPF subjects.

| Sample ID | Status | Lobe | Anatomical Region | Age | Sex | Race | Smoking history | Protocol | Passage |
| --- | --- | --- | --- | --- | --- | --- | --- | --- | --- |
| A160 | Age-matched Control | Lower | Parenchyma | 60 | Male | White | Yes | WLCS | 1 Initial |
| A160 | Age-matched Control | Lower | Parenchyma | 60 | Male | White | Yes | WLCS | 6 Final |
| A160 | Age-matched Control | Lower | Parenchyma | 60 | Male | White | Yes | Negative Fraction | 1 Initial |
| A160 | Age-matched Control | Lower | Parenchyma | 60 | Male | White | Yes | Negative Fraction | 6 Final |
| A001 | Age-matched Control | Lower | Parenchyma | 66 | Female | White | No | Outgrowth | 3 Initial |
| A001 | Age-matched Control | Lower | Parenchyma | 66 | Female | White | No | Outgrowth | 6 Final |
| U197 | IPF | Upper | Parenchyma | 69 | Male | White | No | WLCS | 1 Initial |
| U197 | IPF | Upper | Parenchyma | 69 | Male | White | No | WLCS | 6 Final |
| U197 | IPF | Upper | Parenchyma | 69 | Male | White | No | Negative Fraction | 1 Initial |
| U197 | IPF | Upper | Parenchyma | 69 | Male | White | No | Negative Fraction | 6 Final |
| U001 | IPF | Upper | Parenchyma | 70 | Male | White | Yes | Outgrowth | 3 Initial |
| U001 | IPF | Upper | Parenchyma | 70 | Male | White | Yes | Outgrowth | 6 Final |
| L197 | IPF | Lower | Parenchyma | 69 | Male | White | No | WLCS | 1 Initial |
| L197 | IPF | Lower | Parenchyma | 69 | Male | White | No | WLCS | 6 Final |
| L197 | IPF | Lower | Parenchyma | 69 | Male | White | No | Negative Fraction | 1 Initial |
| L197 | IPF | Lower | Parenchyma | 69 | Male | White | No | Negative Fraction | 6 Final |

**Supplementary Table 3.** Genes Shared Among Lung Tissue Fibroblasts and Cultured Fibroblasts at Initial and Final Passages.

| Gene | Category |  | Function/Description |
| --- | --- | --- | --- |
| COL1A1 | ECM Structure | Extracellular matrix proteins | Collagen type I alpha 1 chain; major component of extracellular matrix, essential for tissue structure and fibrosis |
| COL5A1 | ECM Structure | Extracellular matrix proteins | Collagen type V alpha 1 chain; regulates collagen fibril assembly and tissue organization |
| CYGB | Metabolism | Metabolic enzymes | Cytoglobin; oxygen sensor and regulator of fibrosis, involved in oxidative stress response |
| DLL1 | Cell Signaling | Signal transduction pathways | Delta-like 1; Notch signaling ligand, regulates cell fate and differentiation |
| DTX4 | Cell Signaling | Signal transduction pathways | Deltex E3 ubiquitin ligase 4; regulates Notch signaling pathway |
| ECHDC3 | Metabolism | Metabolic enzymes | Enoyl-CoA hydratase domain containing 3; involved in fatty acid metabolism |
| ECM2 | ECM Structure | Extracellular matrix proteins | Extracellular matrix protein 2; regulates ECM organization and cell adhesion |
| EDNRA | Cell Signaling | Signal transduction pathways | Endothelin receptor type A; mediates vasoconstriction and fibroblast activation |
| EGR1 | Transcription Factor | Gene expression regulation | Early growth response 1; immediate-early gene regulating cellular growth and differentiation |
| EPHX1 | Metabolism | Metabolic enzymes | Epoxide hydrolase 1; detoxification enzyme, metabolizes reactive epoxides |
| EXT1 | ECM Modification | Extracellular matrix proteins | Exostosin glycosyltransferase 1; synthesizes heparan sulfate, important for ECM organization |
| F11R | **Cell Adhesion** | **Cell-cell interactions** | F11 receptor (JAM-A); junctional adhesion molecule, regulates tight junctions and migration |
| FAM20A | **Cell Signaling** | **Signal transduction pathways** | Family with sequence similarity 20 member A; pseudokinase involved in biomineralization |
| FAM43A | **Cell Signaling** | **Signal transduction pathways** | Family with sequence similarity 43 member A; involved in actin cytoskeleton regulation |
| FBLN5 | **ECM Structure** | **Extracellular matrix proteins** | Fibulin 5; elastin-binding protein, essential for elastic fiber assembly and vascular integrity |
| FHL1 | **Cell Signaling** | **Signal transduction pathways** | Four and a half LIM domains 1; mechanosensor involved in muscle development and signaling |
| FIBIN | **Development** | **Tissue development** | Fin bud initiation factor; regulates appendage development and tissue morphogenesis |
| FMO2 | **Metabolism** | **Metabolic enzymes** | Flavin-containing monooxygenase 2; xenobiotic metabolism and oxidative stress response |
| FOS | **Transcription Factor** | **Gene expression regulation** | Fos proto-oncogene; component of AP-1 complex, regulates proliferation and differentiation |
| FRAS1 | **ECM Structure** | **Extracellular matrix proteins** | Fraser extracellular matrix complex subunit 1; important for basement membrane integrity |
| FZD4 | **Cell Signaling** | **Signal transduction pathways** | Frizzled-4; Wnt receptor involved in angiogenesis and vascular development |
| GABRE | **Cell Signaling** | **Signal transduction pathways** | GABA A receptor epsilon; neurotransmitter receptor, may have non-neuronal roles |
| GDF10 | **Cell Signaling** | **Signal transduction pathways** | Growth differentiation factor 10; TGF-beta superfamily member, regulates tissue development |
| INMT | **Metabolism** | **Metabolic enzymes** | Indolethylamine N-methyltransferase; involved in tryptamine and melatonin metabolism |

